# Discrimination of spectrally sparse complex-tone triads in cochlear implant listeners

**DOI:** 10.64898/2026.03.20.712905

**Authors:** Marie-Luise Augsten, Martin J. Lindenbeck, Bernhard Laback

**Affiliations:** Acoustics Research Institute, Austrian Academy of Sciences, Dominikanerbastei 16, 1010 Vienna, Austria

**Keywords:** cochlear implants, chord discrimination, temporal pitch, place pitch, beating

## Abstract

Cochlear implant (CI) users typically experience difficulties perceiving musical harmony due to a restricted spectro-temporal resolution at the electrode-nerve interface, resulting in limited pitch perception. We investigated how stimulus parameters affect discrimination of complex-tone triads (three-voice chords), aiming to identify conditions that maximize perceptual sensitivity. Six post-lingually deafened CI listeners completed a same/different task with harmonic complex tones, while spectral complexity, voice(s) containing a pitch change, and temporal synchrony (simultaneous vs. sequential triad presentation) were manipulated. CI listeners discriminated harmonically relevant one-semitone pitch changes within triads when spectral complexity was reduced to three or five components per voice, with significantly better performance for three-component compared to nine-component tones. Sensitivity was observed for pitch changes in the high voice or in both high and low voices, but not for changes in only the low voice. Single-voice sensitivity predicted simultaneous-triad sensitivity when controlling for spectral complexity and voice with pitch change. Contrary to expectations, sequential triad presentation did not improve discrimination. An analysis of processor pulse patterns suggests that difference-frequency cues encoded in the temporal envelope rather than place-of-excitation cues underlie perceptual triad sensitivity. These findings support reducing spectral complexity to enhance chord discrimination for CI users based on temporal cues.

## I. INTRODUCTION

Cochlear implants (CIs) are auditory prostheses applied in cases of severe-to-profound cochlear hearing loss or even deafness. An electrode array is implanted into the cochlea and directly stimulates the remaining spiral ganglion neurons of the auditory nerve. Acoustic input is converted by a speech processor into pulsatile stimulation patterns which in turn are transmitted to the electrode array (e.g., Wilson & Dorman, 2008). Although the majority of post-lingually deafened CI listeners can successfully understand speech in quiet listening conditions (Gifford et al., 2008; Wilson et al., 1991), satisfying music perception still remains challenging (for reviews, see Drennan & Rubinstein, 2008; Limb & Rubinstein, 2012; Prevoteau et al., 2018).

Whereas perception of rhythm is largely restored in CI listening, pitch, harmony, and timbre perception are severely degraded for many CI listeners (Brockmeier et al., 2011; Bruns et al., 2016; Limb & Roy, 2014; McDermott, 2004). Of the latter, harmony is the most complex musical feature: It is usually defined as the combination of simultaneous notes (i.e., musical voices) to chords and, further, the combination of chords to chord progressions (Dahlhaus, 1980). Chords or chord progressions convey musical structure, evoke perceptions of consonance/dissonance, and, thus, contribute greatly to the emotional effect of music (e.g., Koelsch et al., 2019; Parncutt, 1989; Steinbeis et al., 2006). Due to the restricted spectral resolution in CI listeners, chord perception is even more challenging than single-pitch or melody perception. For example, similar to speech perception in noise, melodic contour identification in CI listeners is worse when a competing instrument is present (Galvin et al., 2009). To pave the way for understanding and improving harmony perception in CI listeners, the present study examines stimulus parameters that optimize the salience of harmony-related pitch cues within chords for CI listeners.

Several technical and physical constraints of the CI are considered to particularly degrade musical features such as pitch and harmony. First, whereas in a healthy cochlea, approximately 3,500 inner hair cells convert mechanical vibrations to spiral-ganglion neuron activity, with current CIs, only between 12 and 24 electrodes fulfill this function. While the electrode array allows to roughly reproduce the tonotopic stimulation pattern of the acoustically stimulated cochlea, very few distinct stimulation sites, that is, electrode positions, are available for conveying pitch based on the place of stimulation along the basilar membrane (i.e., place cue). Furthermore, most salient pitch perception is assumed to depend on an accurate frequency-place map within the cochlea (Oxenham et al., 2004). Frequently occurring misalignments of frequency bands versus stimulation electrodes in CI listeners result in suboptimal neuronal responses on specific frequencies (Landsberger et al., 2015; Zeng et al., 2014). However, it was shown that normal-hearing (NH) listeners reliably discriminate musically relevant pitch changes of one semitone (ST) within harmonic-complex triads (i.e., three-tone chords) without place cues present in the signal, suggesting that purely temporal mechanisms (see below) may allow functional pitch perception in the context of triads (Graves & Oxenham, 2019). Thus, constrained place cues may not fully prevent CI listeners from discriminating chords in a musically meaningful way.

Second, besides the limited number of electrodes *per se*, one major limitation of CIs are channel interactions, that is, the spread of electric fields across the array and, thus, crude representation of the stimulus spectrum (e.g., Fu & Nogaki, 2005; Laback et al., 2004), which is mainly due to imperfect coupling of the electrode contact with the spiral ganglion neurons of the auditory nerve. This lack of channel independence across sites is considered generally unavoidable (e.g., Wilson & Dorman, 2008).

Third, extracting pitch by auditory neurons entraining their response to temporal features of the incoming acoustic signal across time (i.e., temporal cue) is partly sub-optimal with CIs. For signal transmission, most clinical CI systems use electrical pulse trains with constant rates of 900 to 1500 pulses per second (pps) and electrode which convey sound information only by modulating the amplitude of electric pulses based on the acoustic temporal envelope and not their timing based on the acoustic temporal fine structure. For music instruments and voiced speech, the amplitude modulation encodes the fundamental frequency (F0). Such high-rate envelope-coding strategies, like the continuous interleaved sampling (CIS) approach, achieve high speech perception scores in quiet (e.g., Wilson et al., 1991). Yet, due to their lack of temporal fine-structure cues, they do not provide particularly salient pitch cues. As temporal fine structure is assumed to be especially important for conveying salient pitch in normal hearing (Smith et al., 2002), some stimulation strategies such as FAST (Smith, 2010), FSP (Hochmair et al., 2006), or FS4 (e.g., Riss et al., 2014) aim at conveying temporal fine-structure features of acoustic signals via the pulse timing, resulting in aperiodic lower-rate pulse trains on a few apical electrodes (transmitting the electric fine structure, hereafter referred to as FS). Importantly, despite the FS being derived from properties of the acoustic fine structure, it is processed differently (Müller et al., 2022). In fact, FS sensitivity is more similar to or even better than temporal-envelope sensitivity with large modulation depths (Goldsworthy et al., 2021, 2022; Schnupp et al., 2025), thus, probably reflecting a less distorted and more salient representation of envelope information, but still not comparable to “true” fine-structure information. The benefit of FS processing for music is supported by Looi et al. (2011) who reported higher musical sound quality ratings for FSP compared to CIS. In contrast, however, the perception of beats (i.e., temporal interactions of amplitude modulations within one peripheral filter can even be better in CI compared to normal hearing. For example, CI listeners have been shown to be able to precisely tune a guitar only by the reliance on beating cues (Lu et al., 2014). Also, the detection of amplitude modulation depth, a prerequisite for modulation frequency detection, is enhanced in CI listeners compared to NH listeners up to 300 Hz (Shannon, 1992).

Fourth, CI listeners’ ability to reliably discriminate temporal pitch is limited to rates lower than approximately 400 pps (De Groote et al., 2024; Ihlefeld et al., 2015; Kong et al., 2009; Zeng, 2002), such that at higher rates a rate increase does not result in increased pitch. At higher rates, place pitch is usually more dominant than temporal pitch (e.g., Zeng, 2002). As the number of tonotopically organized stimulation sites in the cochlea (i.e., electrodes) is low, a change in frequency does not necessarily result in a precise change in excitation pattern. However, a rate-place covariation is assumed to be necessary for high-level pitch perception (Carlyon & Deeks, 2002; Oxenham et al., 2004). Finally, the above-mentioned spread of electrical field in electric stimulation results in spectro-temporal interactions, so-called channel interactions (e.g., Boulet et al., 2016), which in turn degrade the temporal coding of pitch both in the FS and in the envelope.

As a consequence, perception of pitch and harmony is generally poor in CI listeners compared to NH individuals. However, there is high variability among CI listeners. This variability is assumed to result from factors such as residual acoustic hearing (Gfeller et al., 2006), pre- or post-lingual deafness (Bruns et al., 2016), accuracy of frequency-place mapping within the cochlea (Di Nardo et al., 2011), processing strategy (Roy et al., 2015), musical training before and after implantation (Gfeller et al., 2008; Shukor et al., 2021), age (Zimmer et al., 2019), and cognitive capacities (Gfeller et al., 2008). Furthermore, performance strongly depends on the physical structure of the acoustic stimulus. Whereas for complex sounds, most NH individuals detect pitch changes of one semitone (ST) or less, pitch detection thresholds in CI listeners range from one to eight STs (Kang et al., 2009) or even up to two octaves (Gfeller et al., 2002). However, with low-rate stimuli directly transmitted to one electrode, rate-pitch discrimination sensitivity often reaches an order corresponding to one ST (e.g., Goldsworthy et al., 2022; Moore & Carlyon, 2005), depending on the training state (e.g., Goldsworthy & Shannon, 2014).

For the discrimination of subtle chord changes, CI performance was found to be low (Brockmeier et al., 2011; Lam et al., 2022) while discrimination performance of larger chord changes (i.e., varying number of chord voices and/or relatively large pitch differences between the chords of a trial), was moderate to high (but highly variable across listeners and depending on the task) with complex or natural sounds (Boeckmann-Barthel et al., 2013; Lam et al., 2022). Zimmer et al. (2019) observed significant discrimination of major, minor, augmented, and diminished chords in seven out of twelve CI listeners while 22 out of 24 NH listeners showed significant sensitivity. It was argued that the degraded perception of harmony in CI listeners may be due to perceptual fusion of multiple pitches into one pitch (Donnelly et al., 2009). The authors investigated CI and NH listeners’ ability to judge the number of perceived simultaneous voices and CI listeners performed at chance level in discriminating two- and three-voice stimuli. Notably, even NH listeners exhibited a bias toward identifying all stimuli as one-voice conditions, although they did less so than CI listeners (see also Huron, 1989).

Most studies investigating harmony perception with CIs used real-world music stimuli (e.g., Brockmeier et al., 2011; Caldwell et al., 2016; Koelsch et al., 2004; Lam et al., 2022; Tillmann et al., 2019) that typically allow only very limited control of relevant stimulus features. Although such investigations are undoubtedly valuable, real-world music stimuli (such as instruments sounds) consist of tones with a large spectral bandwidth, thus, involving a high chance of overloading the narrow spectral and temporal information channel of a CI recipient: Due to spectro-temporal channel interactions, the large number of frequency components potentially degrades both temporal and place pitch cues as auditory-nerve response phenomena such as refractoriness and summation (cf. Boulet et al., 2016) may smear both the effective timing and place of stimulation due to interference from surrounding pulses (emitted from neighboring electrodes). Moreover, as a result of using envelope-based stimulation strategies, the stimuli used in those studies disregard FS-based pitch cues and convey only envelope-based F0 cues (for a comparison of FS and envelope-based pitch, see, e.g., Lindenbeck et al., 2020). Hence, they deliver a less salient type of temporal pitch cue (e.g., Drennan & Rubinstein, 2008; Goldsworthy et al., 2022). Thus, F0 discrimination with such complex sounds is particularly challenging for CI listeners. Observations from these real-world-stimulus studies, which were interpreted as evidence for poor harmony perception, are potentially affected by technical and physiological constraints of electric stimulation such that general conclusions regarding harmony perception with CIs *per se* might not be drawn. We argue that specifically controlling stimulus parameters by using artificial harmonic complex (HC) tones with carefully selected parameters is preferable for investigating basic questions in harmony perception with CIs. HC tones are freely parameterizable, and, unlike natural sounds, can be controlled in the number, frequency, and phase of spectral components, allowing to stimulate selected electrode ranges with desired periodicities. This provides the potential to identify stimulus parameters that lead to more distinct and salient percepts in CI listeners, as supported by Galvin & Fu (2011) who reported best performance in a melody perception task for bandpass-filtered piano tones containing only lower-order harmonics compared to, for example, piano tones with full spectrum. Other studies using controlled music stimuli also found that CI listeners prefer low spectral complexity (Gauer et al., 2019; Nagathil et al., 2017; Nemer et al., 2017).

In addition to using synthetic HC stimuli for control of spectral complexity, we addressed the above-mentioned CI-specific rate limit of about 400 pps by employing F0s up to 320 Hz only. Altogether, compared to natural stimuli, these means provide the potential to increase the salience of place and temporal pitch cues and to reduce channel interactions, particularly for multi-voice conditions.

In the present study, we investigated several stimulus parameters in a three-voice chord (hereafter referred to as triad) same/different discrimination task. The first parameter was the *number of frequency components per voice*. We hypothesized that sparsity of acoustic input increases the salience of place and temporal cues for CI listeners. Thus, the lowest number of components was assumed to yield the best performance.

The second parameter was the *voice(s) in which a semitone (ST) change occurred* (hereafter referred to as VST): In each trial, from one triad to the other, a pitch change occurred either only in the highest voice (“high”), only in the lowest voice (“low”), or in both the lowest and the highest voice (“low-high”). We hypothesized that the conditions high as well as low-high yield better performance than the low condition. We argue that changes in two voices lead to best performance simply due to the amount of changing information compared to a change in only one voice. Furthermore, an advantage of a high-voice F0 change compared to a low-voice F0 change is likely to occur due to a common perceptual bias in processing the highest voice (Crawley et al., 2002; Fujioka et al., 2005; Palmer & Holleran, 1994). We did not vary the middle-voice F0 as it was shown that NH listeners experience difficulties in processing inner voices (Graves & Oxenham, 2019; Huron, 1989), and we observed in pilot tests with one CI listener that middle-voice discrimination was impossible.

The third parameter was the *temporal synchrony* of voice presentation within a triad, that is, voices were either presented simultaneously or sequentially. The more voices are added up simultaneously, the larger the spectro-temporal complexity and, therefore, the larger the potential for channel interactions and temporal (modulation) patterns too complex to yield perceptual sensitivity. One technique for unraveling spectral complexity is to present the voices of a chord successively rather than simultaneously, much like the compositional style of arpeggio or broken chords often used in music. Relying on the well-known phenomenon of the integration of sequential pitches into harmony (cf. Tramo et al., 2001), we expected this technique to largely reduce spectral overlap and, thus, to enhance triad discrimination for CI listeners. To control for potential effects of temporal masking we also varied the duty cycle, that is, the relation of tone vs silence durations within sequential presentation of triads: We tested duty cycles of 50%, 75% and 100%.

### II. PSYCHOPHYSICAL EXPERIMENT

The triad parameters described above were studied in one experiment that involved six post-lingually deafened CI listeners. To justify our hypotheses and particularly to verify our assumption that sequentially presented triads can be well discriminated, we additionally pilot-tested four NH listeners. Methods and results of these pilot tests are summarized in Supplementary A.

### A. Methods

#### 3. Listeners

Six adult post-lingually deafened CI listeners (3 females, 3 males, *M*[*SD*]_age_ = 57.2[14.2]) participated in the experiment. A large number of trial repetitions, as employed here, compensates for the restricted number of listeners. Note that such small-*N* designs are a powerful tool to detect systematic relationships manifesting at the listener level (Smith & Little, 2018). Bilateral CI listeners were tested on their preferred or, based on previous studies, more sensitive ear. A contralateral CI processor or hearing aid was always removed. Demographic and etiological details of the CI listeners are listed in Table 1. As we did not attempt to investigate the impact of possible physiological or technical listener differences on perception and given the small-*N* design, we aimed at recruiting a CI sample as homogenous as possible regarding implant manufacturer, stimulation strategy, insertion depth, residual hearing and onset of deafness to avoid idiosyncratic effects in this realm. In this regard, all CI users were post-lingually deafened in adulthood, used 12-channel implants manufactured by MED-EL GmbH (Innsbruck, AUT) with FSP or FS4 strategy, had at least 28 mm array length, and no residual hearing on the tested ear. Furthermore, participants were recruited based on whether they had decent speech perception (not formally tested) and pitch discrimination abilities (observed in previous studies). Regarding musical training, the sample was balanced such that three of the CI listeners had at least amateur-level musical training with their CI(s) (defined as more than five years of formal instrument training and at least partially identifying as musicians) while three had little to no musical training (defined as five or fewer years of formal instrument training and identifying as non-musicians). Musical sophistication was assessed using an excerpt of the German version of the Gold-MSI (Schaal et al., 2014; for details see Supplementary B). Results are summarized in Table 2. In the experiments, we followed the European Charter of Fundamental Rights, worked along the guidelines of “Good Scientific Practice,” and fulfilled the ethical principles for research involving human subjects. The research protocol was one of the ethics-approved standard protocols for CI research at our laboratory. Participants consented in written form and were paid an hourly wage.

**Table 1.**
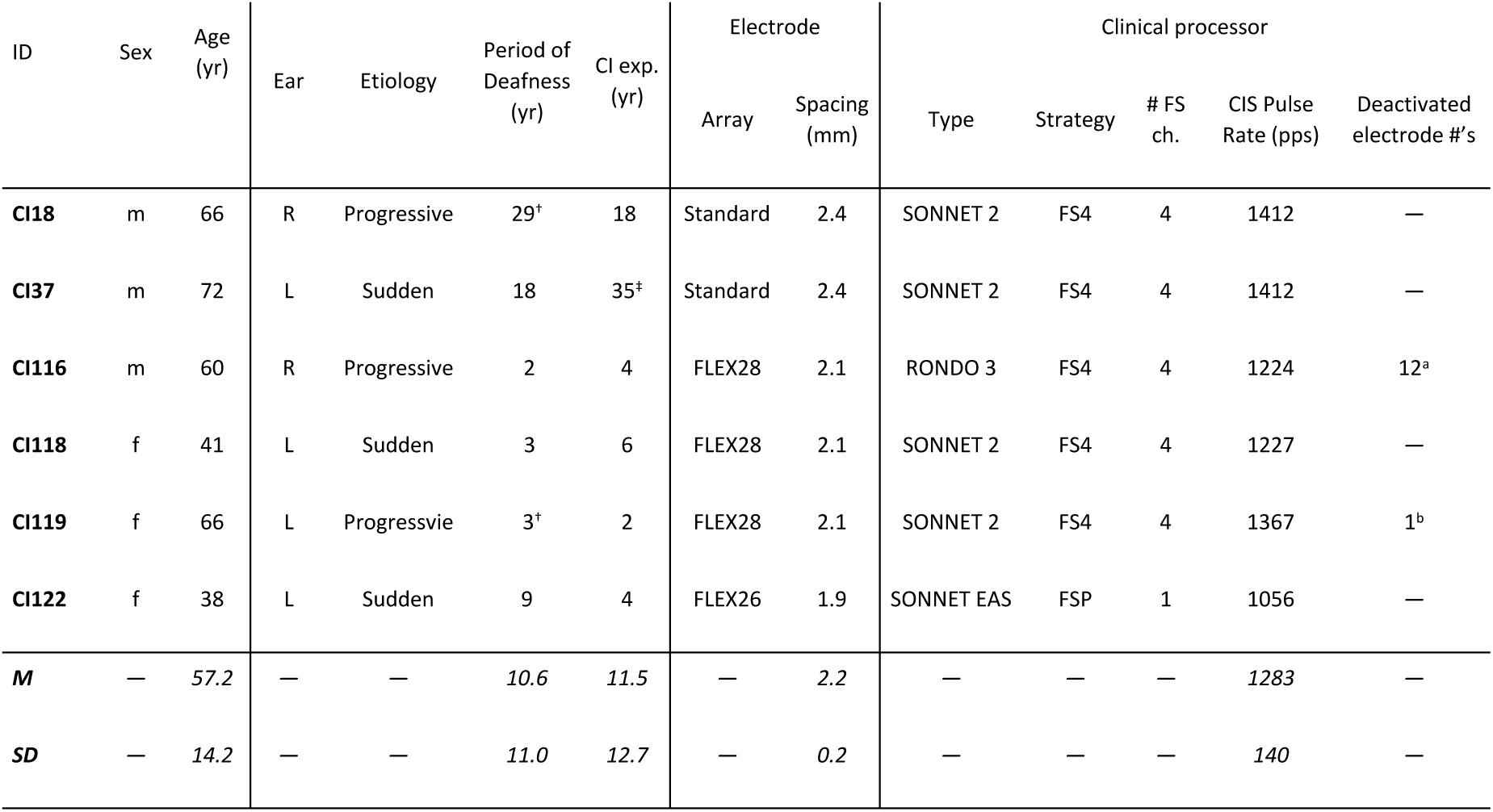
Clinical details for the participating CI listeners. Electrodes numbered from apex to base. Period of deafness refers to years of deafness prior to implantation. Abbreviations: yr…years; FS…fine structure; CIS…continuous interleaved sampling; pps…pulses per second.

**Table 2.**
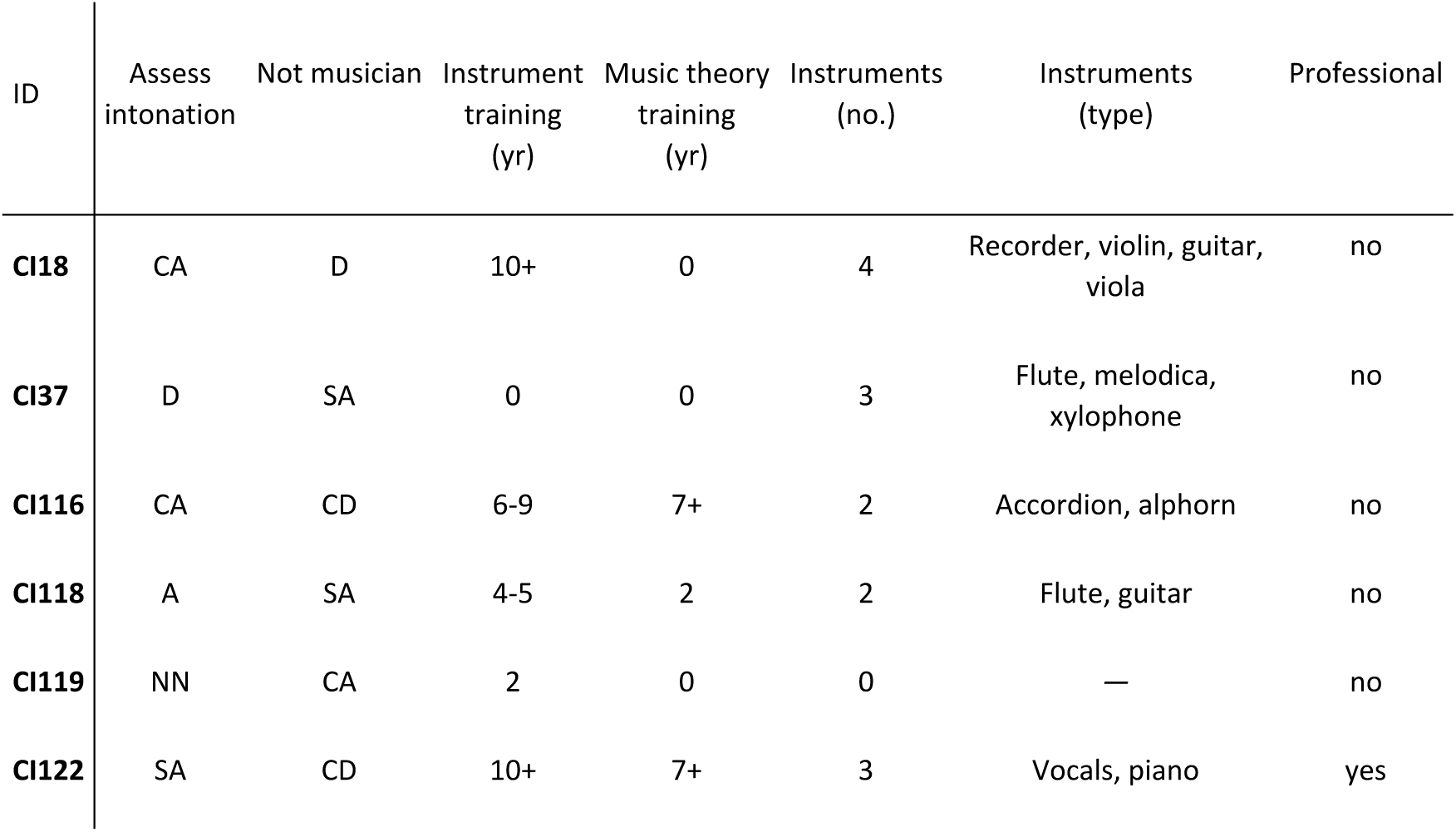
Assessment of CI listeners’ musical sophistication using selected questions from the Gold-MSI questionnaire. See Supplementary B for full questions. Abbreviations: yr…years; CD…completely disagree; SD…strongly disagree; D…disagree; NN…neither disagree nor agree; A…agree; SA…strongly agree; CA…completely agree. Professionality assessment was based on whether participants had a university degree in at least one music-related discipline.

#### 2. Stimuli and apparatus

Stimuli were composed acoustically as harmonic complex tones (HCs) with the first three, five, or nine components of the harmonic series each. These stimuli will be hereafter referred to as HC3, HC5, and HC9, respectively. All components were presented at equal level and added in sine phase. Each stimulus was 400 ms long and had 20-ms raised-cosine onset and offset ramps. Stimuli were generated digitally at a sampling rate of 48 kHz using MATLAB R2021b (The MathWorks, Inc., 2021), converted to the analog domain with an RME Fireface UC (Audio AG, Haimhausen, GER), and amplified by a sonible hp:1 headphone amplifier (sonible GmbH, Graz, AUT). This signal was fed to the auxiliary input of a commercially available and individually configured SONNET 2 Me1510 CI processor (MED-EL GmbH, Innsbruck, AUT), thus, effectively bypassing the processor microphones. This processor will hereafter be referred to as research processor.

To configure the research processor for each CI listener individually, the full configuration of a listener’s clinical CI processor was read out using MAESTRO 9.0.5 (MED-EL GmbH, Innsbruck, AUT). The research processor configuration was largely based on this clinical configuration to keep the stimulation maximally familiar while standardizing pre-processing to maximize comparability. To this end, we never changed the filterbank properties but set the maplaw to default values. Furthermore, the stimulation strategy FS4 (e.g., Riss et al., 2014) was always used with the research processors. FS4 employs the CIS approach (Wilson et al., 1991) on the mid-to-basal electrodes five to 12 while using FS processing (cf. Hochmair et al., 2006) on the four most apical electrodes. Compared to CIS which only transmits acoustic temporal-envelope cues via amplitude modulation of constant-pulse-rate carriers, FS pulses are triggered by bandpass-signal zero-crossings with amplitudes based on the Hilbert envelope. Consequently, FS processing on apical electrodes yields lower and stimulus-dependent (i.e., aperiodic) pulse rates. We restricted our investigation to the FS4 strategy because, as outlined in the Introduction, FS processing is considered particularly advantageous to convey more salient temporal pitch cues. The filterbank properties as well as the CIS pulse rates of the listener-specific research processor configurations are summarized in Table 3. Schematics of pulse trains generated by both FS and CIS processing in response to a HC stimulus are shown in Figure 1. Five of the six CI listeners also used FS4 in their clinical configurations (cf. Table 1). For these listeners, the rest of the clinical fitting, in particular the absolute thresholds and maximum comfortable levels, were copied to the research processor configuration as is. For the other listener, the clinical FSP fitting was converted to FS4 which likely changed the effective pulse rates on the apical four electrodes. To compensate for this, absolute thresholds and maximum comfortable levels were adjusted on these electrodes using MAESTRO. The configuration of the research processor’s pre-processing stages was identical for all CI listeners. Noise-reduction algorithms and adaptive processing were deactivated. The automatic gain control was set to minimal sensitivity, thus largely deactivating it. This was done to minimize non-linear distortion of the acoustic signal and to avoid idiosyncratic behavior of the research processor during the experiments. The input level was fixed at maximum.

**Figure 1.**
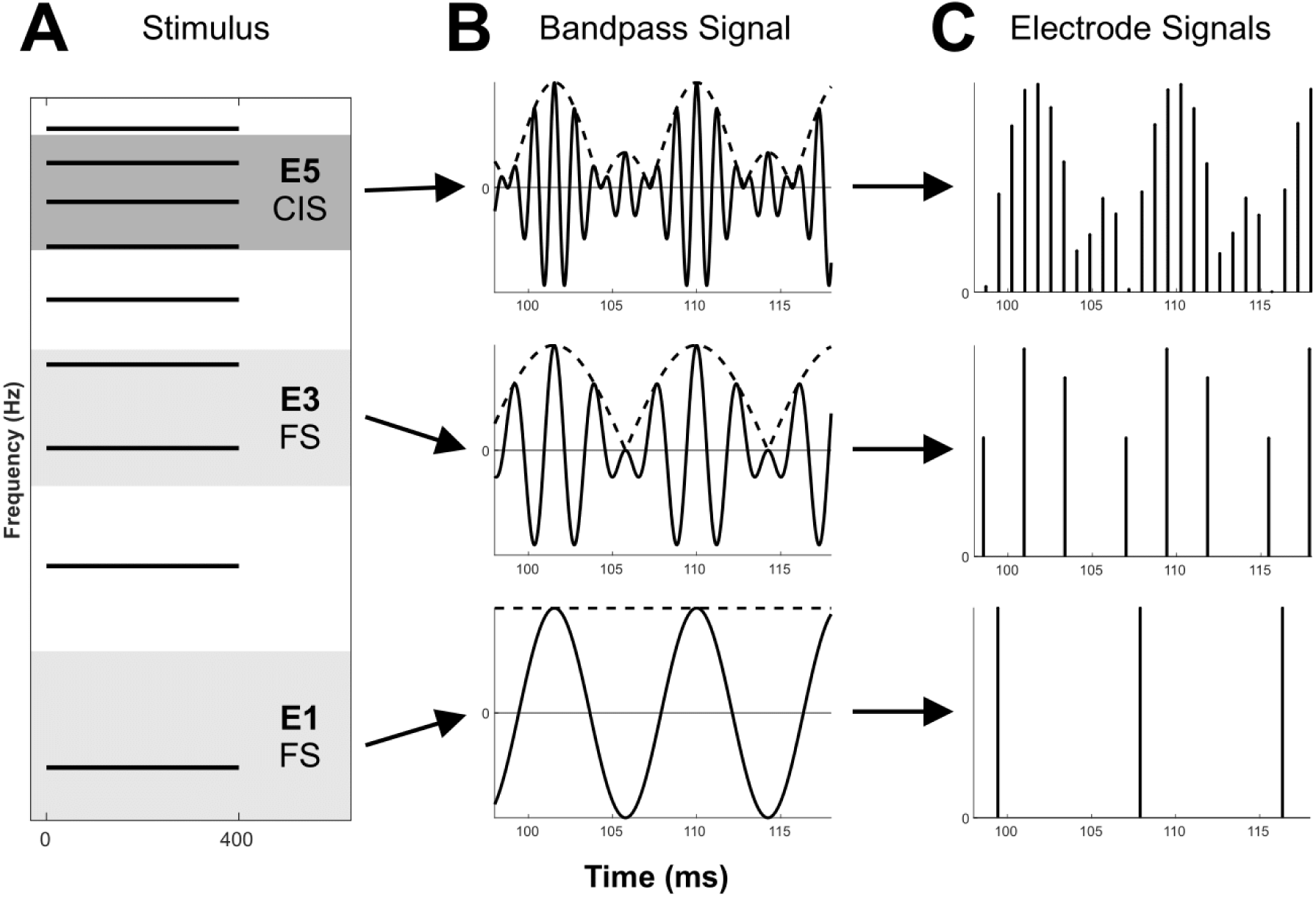
Schematic processing of an HC9 stimulus with an F0 of 118 Hz. **A** Frequency components of the HC stimulus and average passbands (cf. Table 2) for electrodes one, three, and five (E1/E3/E5). FS processing is used on E1 and E3 (light gray shading), CIS processing is used on E5 (dark gray shading). **B** 20-ms snippets of the bandpass signals for E1, E3, and E5 (solid lines) and corresponding Hilbert envelopes (dashed lines). **C** 20-ms snippets of the electrode signals (i.e., pulse trains) after FS or CIS processing (including a logarithmic maplaw). Only positive phases are shown.

**Table 3.** Configuration of the SONNET 2 research processor for the participating CI listeners. Electrodes numbered from apex to base. Electrode numbers in parentheses refer to CI119 who had electrode 1 deactivated in her clinical fitting. “Min.” and “Max.” refer to the 3-dB cut-off frequencies of the respective bandpass filters and together define the filter passbands of FS channels. Abbreviations: CIS…continuous interleaved sampling; pps…pulses per second.

| ID | CIS Pulse Rate (pps) | Deactivated electrode #'s | Electrode 1 (2) |  | Electrode 2 (3) |  | Electrode 3 (4) |  | Electrode 4 (5) |  |
| --- | --- | --- | --- | --- | --- | --- | --- | --- | --- | --- |
|  |  |  | Min. (Hz) | Max. (Hz) | Min. (Hz) | Max. (Hz) | Min. (Hz) | Max. (Hz) | Min. (Hz) | Max. (Hz) |
| **CI18** | 1412 | — | 70 | 225 | 225 | 380 | 380 | 535 | 535 | 690 |
| **CI37<sup>†</sup>** | 1302 | — | 70 | 170 | 170 | 300 | 300 | 469 | 469 | 690 |
| **CI116** | 1224 | 12 | 70 | 205 | 156 | 361 | 288 | 562 | 499 | 780 |
| **CI118** | 1227 | — | 70 | 170 | 170 | 300 | 300 | 469 | 469 | 690 |
| **CI119** | 1367 | 1 | 70 | 181 | 181 | 327 | 327 | 523 | 523 | 787 |
| **CI122<sup>†</sup>** | 1250 | — | 70 | 170 | 170 | 300 | 300 | 469 | 469 | 690 |
| **M** | 1297 | — | 70 | 187 | 179 | 328 | 316 | 505 | 494 | 721 |
| **SD** | 78 | — | 0 | 23 | 24 | 35 | 34 | 41 | 30 | 48 |

<sup>†</sup> Absolute threshold and maximum comfortable level of the fine-structure electrodes (1 to 4) were checked and adjusted if necessary.

#### 3. Procedure

##### a Loudness balancing of single voices

Listeners performed adaptive loudness balancing of single HCs using the weighted up-down staircase procedure (Kaernbach, 1991). Because pilot tests showed only negligible effects of the number of HC components, only HC5 stimuli were balanced.

First, the loudness of a HC5 reference stimulus with an F0 of 196.0 Hz (piano tone G3) was informally set to a comfortable-to-loud level. Second, four target HC5 stimuli with F0s of 82.4, 123.5, 293.7, and 440 Hz (piano tones E2, B2, D4, and A4, respectively) were formally balanced in loudness to the reference. To this end, reference and target stimulus were presented in randomized order separated by a silent gap of 500 ms and listeners were asked which of the two stimuli was louder (two-interval, two-alternative, forced-choice task). They responded on a conventional computer keyboard by pressing F for ‘first stimulus louder’ and J for ‘second stimulus louder’.

To estimate for each target stimulus the 50% point on the psychometric function of loudness (i.e., the point of subjective equality), two blocks of two staircases each were run, resulting in a total of 16 staircases per listener. In each block, one staircase converged at the 75% point on the psychometric function while the other staircase converged on the 25%. Each staircase started with the target stimulus being 20 dB louder (75% staircase) or softer (25% staircase) than the reference, respectively, and was terminated after eight turnarounds. Start level roving of ±4 dB was applied (uniform distribution). For two participants (CI118 and CI119), due to loudness constraints, the start level was decreased to 10±2 dB. The initial geometric mean step size (cf. Kaernbach, 1991) was 4.0 dB and was decreased after four turnarounds to 2.9 dB. The last four turnarounds of the two staircases were arithmetically averaged as estimates of balanced loudness. In total, eight such estimates were collected. The loudness balancing was implemented in PsychoPy v2021.2.3 (Peirce et al., 2019) and based on an existing Python implementation of Kaernbach’s procedure (Mathias, 2013).

The estimates of balanced loudness were fitted with polynomials to get a continuous loudness function between 82.4 and 440 Hz per CI listener. The fit with the highest adjusted R^2^ was selected, hence, protecting against overfitting. Orders of the polynomial fits were between two and four. The fitting was done in MATLAB R2021b (The MathWorks, Inc., 2021).

##### b Pretest: Discrimination of single voices

To estimate basic single-tone (i.e., voice)-discrimination ability with HC stimuli both as a precondition for inclusion in the main test and as a potential predictor of triad-discrimination ability, participants completed a single-voice discrimination test prior to the main triad discrimination test. Listeners were presented with two 400-ms single voices with different F0s and separated by a 500-ms gap. They were asked to respond whether the stimuli were the same or different by pressing M for ‘same’ or V for ‘different’ (same/different task) on a conventional computer keyboard. The proportion of same trials was 50%. For task justification, see Section II.A.3.c). To match F0s to the low and high triad voices in the main experiment, we tested two F0 ranges, one with a nominal F0 of 125.3 Hz (‘low’) and the other with a nominal F0 of 289.5 Hz (‘high’). For each nominal F0, HC3, HC5, and HC9 stimuli were tested and the difference between the lower and higher F0 was either one or two STs (5.9 or 12.2%, respectively). For both F0 differences, the geometric mean of lower and higher F0 matched the nominal F0. For example, for the low nominal F0 of 125.3 Hz and a two-ST F0 difference, the lower F0 was 118.2 Hz and the higher F0 was 132.7 Hz. For each combination of nominal F0, F0 difference, and stimulus type (i.e., number of components per voice), all four possible combinations of lower and higher F0 were presented 25 times each, resulting in 100 repetitions per condition. Across all conditions, the minimum F0 was 118.2 Hz and the maximum F0 was 306.7 Hz. Level roving was not applied. Trial-by-trial feedback was provided. Hits (i.e., same trials followed by a ‘same’ response) and False Alarms (i.e., different trials followed by a ‘same’ response) were subsequently converted to *d’* (Green & Swets, 1966). For this setup, the *d’* range was ±4.1 which corresponds to a hit rate of 98% and a false alarm rate of 2% (i.e., *N*-1 hits and one false alarm to prevent the score from reaching infinity). The pretest consisted of 1200 trials in total. All trials were tested in randomized order in a single block. Listeners were allowed to take breaks at any time. Some initial training trials were allowed for familiarization with the task, if necessary.

##### c Main experiment: Discrimination of simultaneously and sequentially presented triads

The main experiment investigated triad discrimination with a same/different task as in the pretest. We chose this task to allow for a more holistic perception (e.g., of voice interactions, tension, or pleasantness) as opposed to priming the listeners to one specific stimulus dimension (i.e., F0). Such priming is regularly done with a ‘tone height’ task where the listener judges which of two tones is higher in pitch (e.g., Bruns et al., 2016; Ihlefeld et al., 2015; Kang et al., 2009). While tone height is arguably the most relevant stimulus dimension for single-voice discrimination, it may not be the most relevant dimension for triad discrimination as a proxy to “harmonic listening” in everyday life. For example, some studies concluded that either integration of multiple concurrent voices into one perceptual stream is preferred over segregating the voices (Bigand et al., 2000) or both segregation and integration are equally important mechanisms for music perception (Uhlig et al., 2013).

Still, a same/different task is more prone to listeners basing their responses on confounding stimulus dimensions than a lower/higher task. To this end, we identified loudness as the most relevant confounding cue and aimed to minimize its influence on same/different discrimination by balancing the loudness across all occurring F0s. Furthermore, listeners were instructed that loudness is a task-irrelevant stimulus dimension.

Triads were presented in open position (i.e., there was a minimum of seven STs between two triad voices); hence, triad voices covered the range of F0s between 116.5 Hz (piano tone B♭2) and 311.1 Hz (piano tone E♭4). The nominal F0 was 190.4 Hz.

The first experimental parameter was the number of components per triad voice. We used HC3, HC5, and HC9 stimuli. The amplitudes for the three constituting voices were taken from the loudness fittings (cf. Section II.A.3.a)).

The second experimental parameter was the VST. Since we found no sensitivity for ST changes in the middle voice in pilot tests, we only tested ST changes in the low or high voices, or both low and high voice at the same time (i.e., VST conditions, low, high, and low-high). In musical terms, the low condition corresponded to discrimination between a major and an augmented chord in second inversion. The high condition corresponded to discrimination between a major and an augmented chord in first inversion. The low-high condition corresponded to discrimination between a diminished and a major chord in first inversion (see Figure 2). Thus, three different triad pairs were presented. The order of presented triads within a pair was balanced. The third experimental parameter was temporal synchrony. We tested simultaneous triads with HC3, HC5, and HC9 stimuli. An example discrimination trial for a simultaneous HC3 triad is shown in Figure 3. Sequential triads were tested only with HC3 and HC9 due to time constraints. The presentation order of sequential triad voices varied randomly. For sequential triads, we also tested the duty cycle (DC, in %), that is, the proportion between voice duration *T_voice_*and silence duration *T_sil_* (i.e., DC = 100 · *T_voice_*/(*T_voice_* + *T_sil_*)). We tested three DCs of 100, 75, and 50%, respectively. Because we preserved the total triad duration of 400 ms (i.e., 3 · *T_voice_* + 2 · *T_sil_* = 400 ms ∀ DC), voice durations were 133, 109, and 80 ms for DC100, DC75, and DC50, respectively. Accordingly, silence durations were 0, 36, and 80 ms. The DC is illustrated in Figure 4. Because the overall DC effect might be confounded by an effect of the voice duration, we furthermore included simultaneous control conditions with reduced duration of 80 ms (corresponding to the sequential DC50 condition), for HC3 and HC9. To somewhat compensate for the shorter total duration of these control conditions, the triad amplitude was increased by 3.0 dB as compared to the respective 400-ms conditions. Furthermore, the sequential-triad voice amplitudes were increased by 1.2 dB (DC75) and 3.0 dB (DC50), respectively, relative to the (loudness-balanced) single-voice amplitudes.

**Figure 2.**
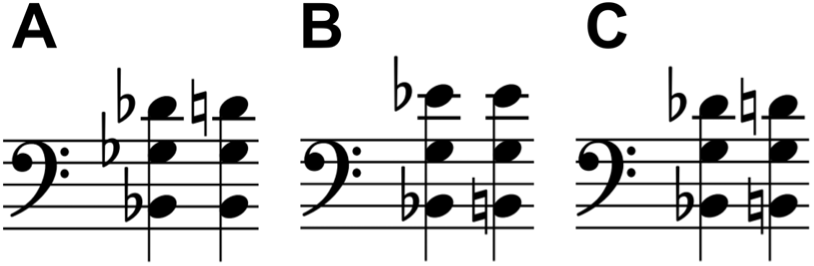
Musical note examples of triad stimuli used in the main experiment with 1-semitone frequency changes per changed voice. **A** Change in the high voice. **B** Change in the low voice. **C** Change in both high and low voice.

**Figure 3.**
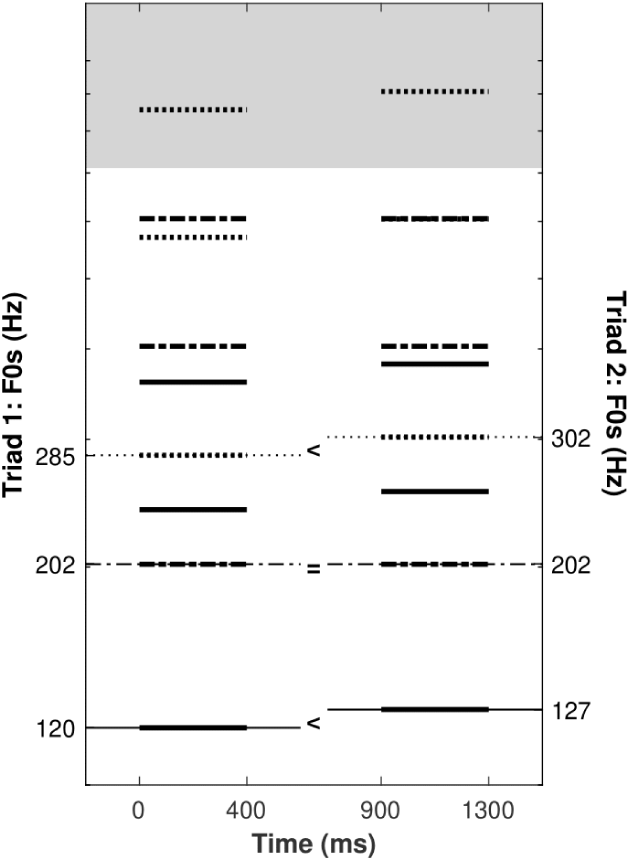
Example discrimination trial for a simultaneous HC3 triad. Low-F0 voice with solid lines, middle-F0 voice with dash-dotted lines, and high-F0 voice with dotted lines. Between the two stimuli, both the low and the high voice differ in F0 by one semitone (i.e., 5.95% in equal temperament).

**Figure 4.**
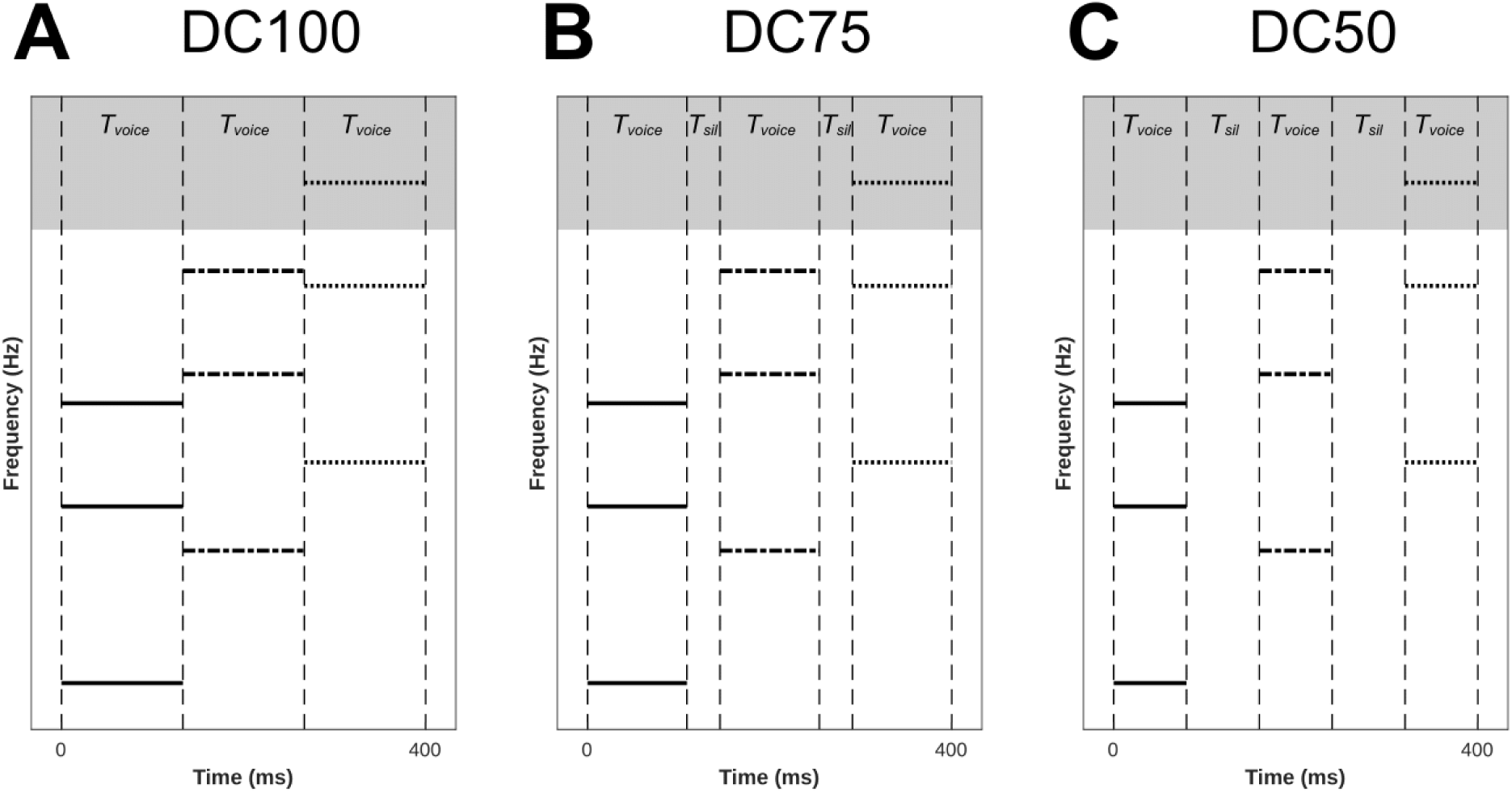
Example sequential triads with different duty cycles (DCs). **A** DC100 (i.e., no silence between voices). **B** DC75 (i.e., *T_voice_* = 3 · *T_sil_*). **C** DC50 (i.e., *T_voice_* = *T_sil_*). The total triad duration is always 400 ms. All other aspects as in Figure 2.

All other aspects of task and procedure were as described for the single-voice pretest (cf. Section II.A.3.b)). The main experiment consisted of 6900 trials in total. For the simultaneous condition, each combination of number of components, VST and duration was repeated 100 times (blocked by duration), resulting in two blocks with 900 (400 ms: HC3, HC5, and HC9) and 600 (80 ms: HC3 and HC9) trials, respectively. Trial repetitions were applied similarly for sequential triads and they were presented block-wise by number of components and DC, resulting in 6 blocks with 900 trials each. Again, breaks were allowed at any time and participants were explicitly and frequently encouraged to take breaks, reducing the likelihood that they experienced fatigue during the tests. Pure test time (i.e., including loudness balancing and pretest but without individual breaks) was approximately 5.5 hours, which was split in two separate test days.

#### 4. Statistical analysis

Due to the identical experimental setup for all CI listeners, we expected large inter-individual differences and *d*’ distributions skewed towards zero, both paving the way for violations of assumptions of conventional repeated-measures analyses of variance. Hence, *d’*s (computed as stated above) were converted to ranks using the aligned-rank transformation and subjected to a mixed-effects analysis of variance (hereafter referred to as ART ANOVA; Wobbrock et al., 2011). This is a rank-based, non-parametric test for factorial experiments, like Friedman ANOVA except that it allows for multiple factors and, crucially, interactions. The significance level was always 5%. To correct for multiple comparisons while preserving power in (two-sided) post-hoc *t*-tests, we controlled the false-discovery rate (Benjamini & Hochberg, 1995) instead of the family-wise error. As effect-size metric, we used the generalized η^2^ (η_G_^2^) (Bakeman, 2005; Lakens, 2013; Olejnik & Algina, 2003), derived from conventional repeated-measures analyses of variance. It is comparable across both within-subjects and between-subjects designs and classifies effect sizes as either small (η_G_^2^ ≥ .02), medium (η_G_^2^ ≥ .13), or large (η_G_^2^ ≥ .26) (Bakeman, 2005).

To assess whether *d*’ scores were significantly larger than zero, we performed bootstrapped one-sided one-sample *t*-tests (Efron & Tibshirani, 1994).

To calculate partial within-subjects correlations *r_p_*(cf. Bland & Altman, 1994, 1995) between single-voice and triad *d’* scores normalized to control for the effects of number of frequency components per voice and the F0 range/VST, we calculated Pearson’s *r* for between single-voice and triad *d’* scores in a linear regression model with triad *d’* as dependent variable and single-voice *d’* as continuous predictor. Furthermore, the categorical predictors CI listener, number of frequency components per voice, F0 range/VST, and the interaction of the latter two, predictors were included in the model. The partial correlation *r_p_* was derived from the model *t* statistics using relation 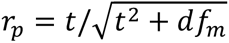 where *t* is test statistic for the continuous predictor single-voice *d’* and *df_m_* are the residual model degrees of freedom (e.g., Fritz et al., 2012). For *r_p_*, 95% confidence intervals are denoted in brackets.

Statistical analyses were conducted with R 4.3.2 (R Core Team, 2023) and the packages ARTool (Kay et al., 2021) and MKinfer (Kohl, 2019/2024).

### Results

#### 1. Pretest

Discrimination sensitivity (in terms of *d*’) is shown in Figure 5 as a function of number of components with F0 difference as parameter and in separate panels per F0 Range. All CI listeners showed sensitivity to single-voice differences in most conditions and were therefore included in the main triad discrimination test. We conducted an ART ANOVA on *d’* scores with number of components (i.e., 3 vs. 5 vs. 9), F0 Range (i.e., low vs. high, separate sub-panels), and F0 difference (FD; i.e., 1 vs. 2 STs) as within-participant variables. We found a main effect of number of components, *F*(2, 10) = 4.85, *p* = .034, η²_G_ = 0.038, a main effect of FD, *F*(1, 5) = 40.48, *p* = .001, η²_G_ = 0.183, and an interaction between number of components and FD, *F*(2, 10) = 6.48, *p* = .016, η²_G_ = 0.023, due to higher performance for HC9 than HC3 triads, *t*(10) = 3.25, *p* = .026, and marginally higher performance for HC5 than HC3 triads, *t*(10) = 2.42, *p* = .054, when FD was 2 STs. To investigate whether *d’* scores were significantly larger than zero, bootstrapped one-sided *t*-tests were calculated for each factor combination. All scores were significantly different from zero (*p* ≤ .0167).

**Figure 5.**
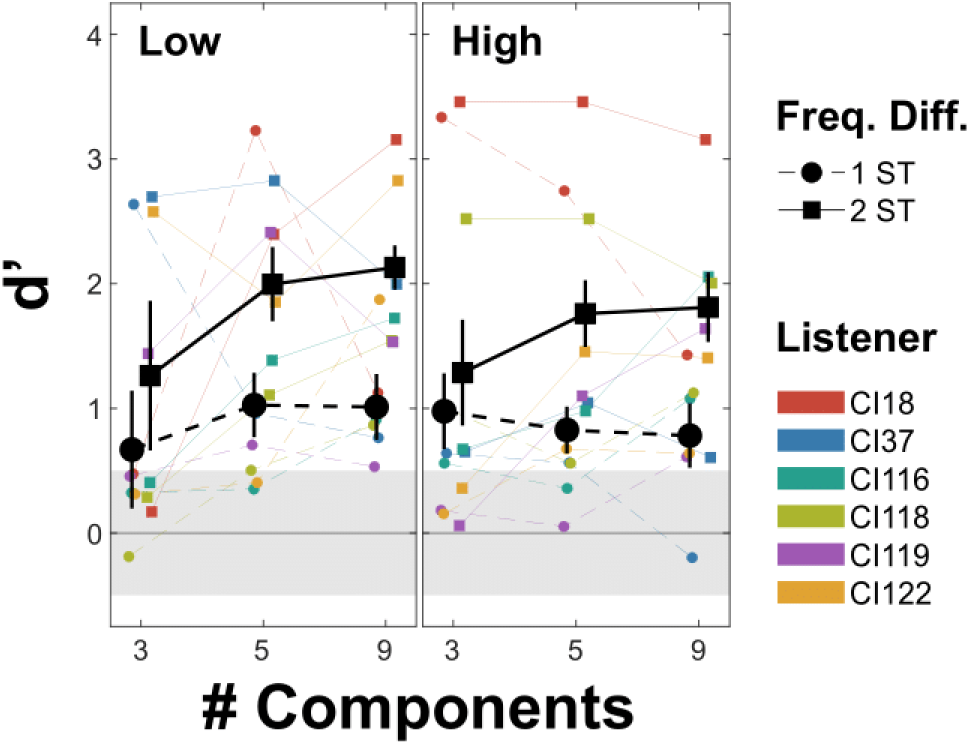
Discrimination sensitivity in terms of *d’* for single voices as function of frequency components per voice. Panels distinguish F0 ranges (low vs. high). Marker shapes and line types denote the frequency difference (FD) to be discriminated in terms of STs. Error bars denote normalized (i.e., within-subjects) standard errors (Cousineau, 2005; Morey, 2008). The grey ribbons indicate chance performance for 100 repetitions. Individual data in color.

#### 2. Main experiment

Analyses were initially split into simultaneous and sequential conditions to reduce the number of independent variables and, thus, to preserve power. Still, selected conditions were subsequently compared directly.

##### a Discrimination of simultaneous triads

Discrimination sensitivity for simultaneous triads is shown in Figure 6 as a function of number of components with VST as parameter. On the group level, sensitivity decreased with increasing number of components and was highest for the VST low-high condition and lowest for the VST low condition.

**Figure 6.**
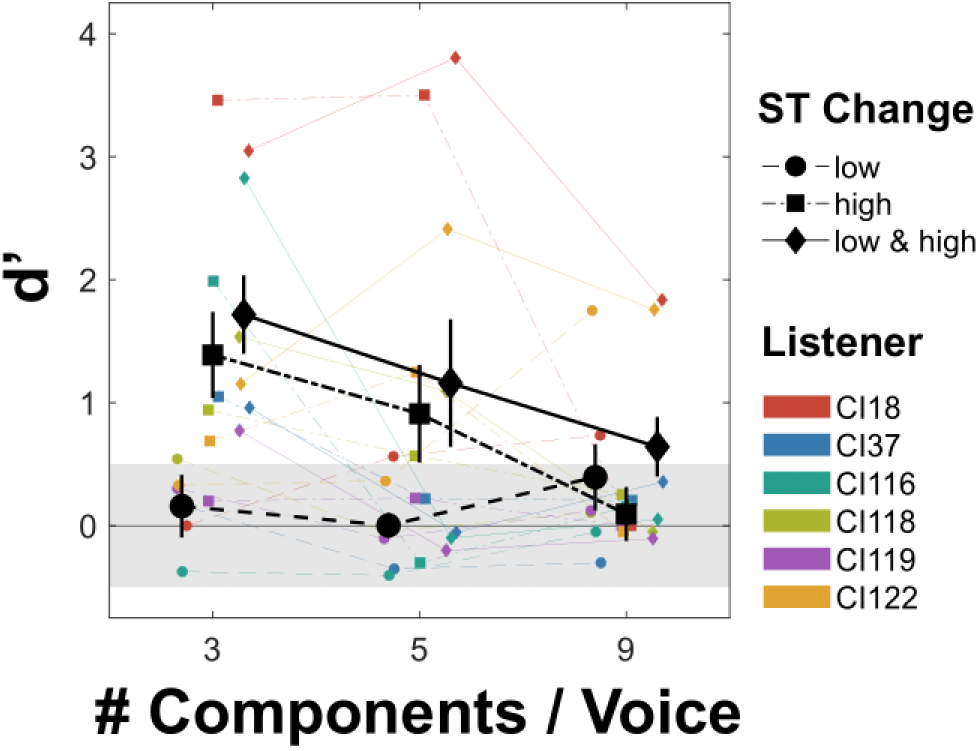
Discrimination sensitivity in terms of *d’* for simultaneous triads as function of frequency components per triad voice. Marker shapes and line types denote the triad voice(s) with a ST change to be discriminated. All other aspects as in Figure 5.

For the analysis of CI listeners’ *d*’ scores on simultaneous-triad discrimination, we ran an ART ANOVA with VST (i.e., low vs. high vs. low-high) and number of components (i.e., 3 vs. 5 vs. 9 components per triad voice) as within-participant variables.

The analysis revealed a main effect of VST, *F*(2, 10) = 11.83, *p* = .002, η²_G_ = 0.173, and an interaction between VST and number of components, *F*(4, 20) = 3.10, *p* = .039, η²_G_ = 0.095. As the interaction resulted from chance-level responses to triads with a low-voice change for all numbers of components, to gain test power for post-hoc testing the number of components hypothesis, we ran another ART ANOVA excluding the VST low condition, leaving only high vs. low-high. This analysis yielded a main effect of number of components, *F*(2, 10) = 5.85, *p* = .021, η²_G_ = 0.179, due to higher performance for HC3 than HC9 triads, *t*(10) = 2.96, *p* = .043.

Again, bootstrapped *t*-tests against zero were calculated for each combination of number of components and VST, showing that HC3 triads were significantly discriminable for the VST conditions low-high, *M* = 1.72, *SE* = 0.35, *t*(5) = 4.28, *p* = .001, and high, *M* = 1.38, *SE* = 0.41, *t*(5) = 2.90, *p* = .003. Furthermore, HC5 triads were significantly discriminable for VST conditions low-high, *M* = 1.16, *SE* = 0.59, *t*(5) = 1.73, *p* = .041, and high, *M* = 0.90, *SE* = 0.47, *t*(5) = 1.63, *p* = .029. Also, for the VST low-high conditions, HC9 triads were significantly discriminable, *M* = 0.64, *SE* = 0.33, *t*(5) = 1.72, *p* = .037. *T*-tests of the other factor combinations were not significant (*p* ≥ .065).

##### b Discrimination of sequential triads

Discrimination sensitivity for sequential triads is shown in Figure 7 for HC3 triads in panel A and for HC9 triads in panel B as a function of DC with VST as parameter. On the group level, sensitivity was mostly absent. On the individual level, CI18 is a notable exception.

**Figure 7.**
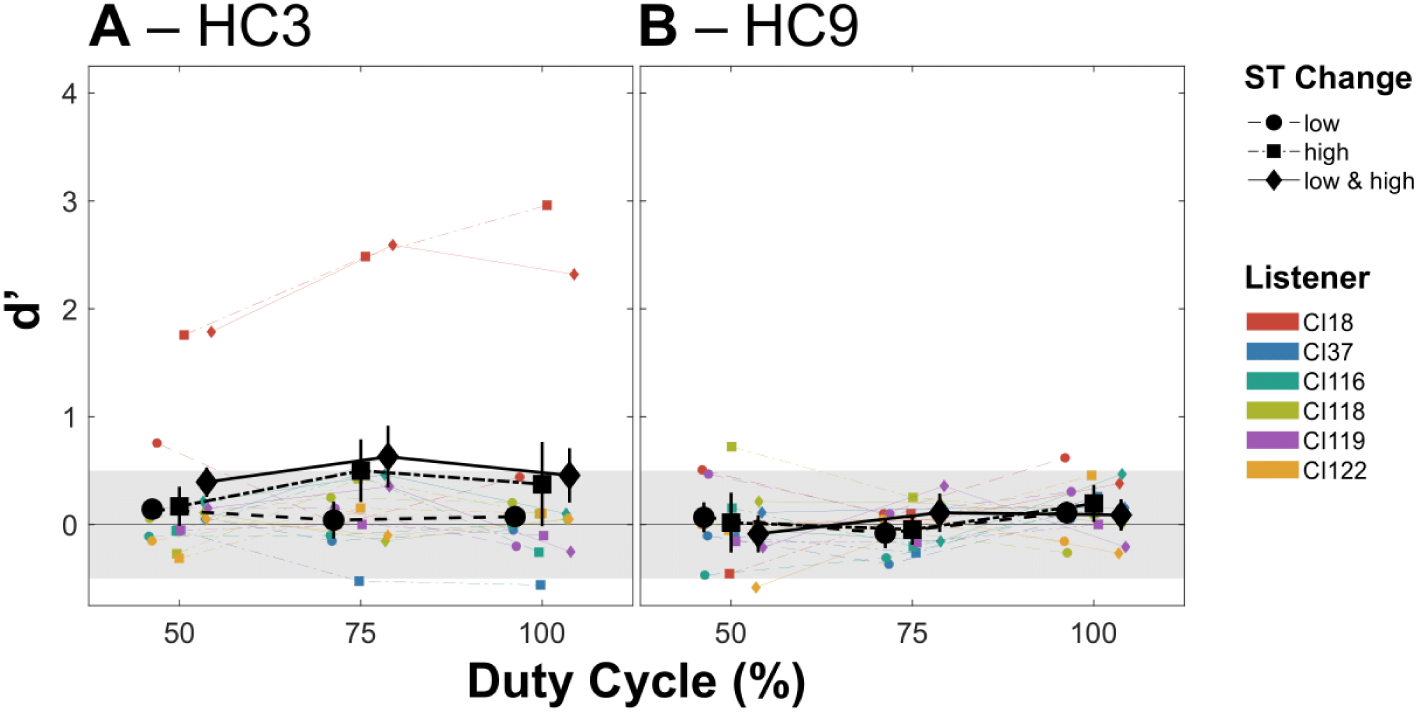
Discrimination sensitivity in terms of *d’* for sequential **A** HC3 and **B** HC9 triads as a function of DC. All other aspects as in Figure 6.

As visual inspection of the sequential-triad discrimination data revealed that HC9 triads yielded no discrimination sensitivity, we ran the analysis for HC3 triads only. We tested two within-subjects variables: VST and DC (i.e., 50 vs. 75 vs. 100%). However, neither the main effects, *F*(2, 10) ≤ 2.61, *p* ≥ .122, η²_G_ ≤ 0.049, nor the interaction, *F*(4, 20) = 1.27, *p* = .313, η²_G_ = 0.011, reached significance.

Nevertheless, sequential HC3 triads were significantly discriminable (i.e., *d’* scores larger than zero) in two conditions, that is, in the VST low-high condition combined with a DC of 50%, *M* = 0.39, *SE* = 0.21, *t*(5) = 1.41, *p* = .048, or combined with a DC of 75%, *M* = 0.63, *SE* = 0.34, *t*(5) = 1.53, *p* = .034. However, these results do not or only barely exceed the population sensitivity threshold (*d’* = 0.5) but instead are skewed by the extraordinarily high performance of CI18.

#### 3. Single-voice sensitivity as a predictor of simultaneous-triad sensitivity

To assess whether CI listeners who perform better in single-voice discrimination also perform better in triad discrimination, that is, whether single-voice sensitivity is predictive of triad sensitivity, we calculated a within-subjects correlation between single-voice *d’* scores and triad *d’* scores that factored out the effects of number of frequency components per voice and F0 range/VST (and their interaction). The correlation between these normalized *d’* scores is shown in Figure 8. At the group level, the correlation was significant, *r_p_*(24) = .54 [.22, .73], *p* = .004, *N* = 36, and thus explained between 5 and 53% of the residual variance (95% confidence interval). Musicians (particularly CI18 and CI122) tended to show stronger correlations than non-musicians (particularly CI37 and CI118).

**Figure 8.**
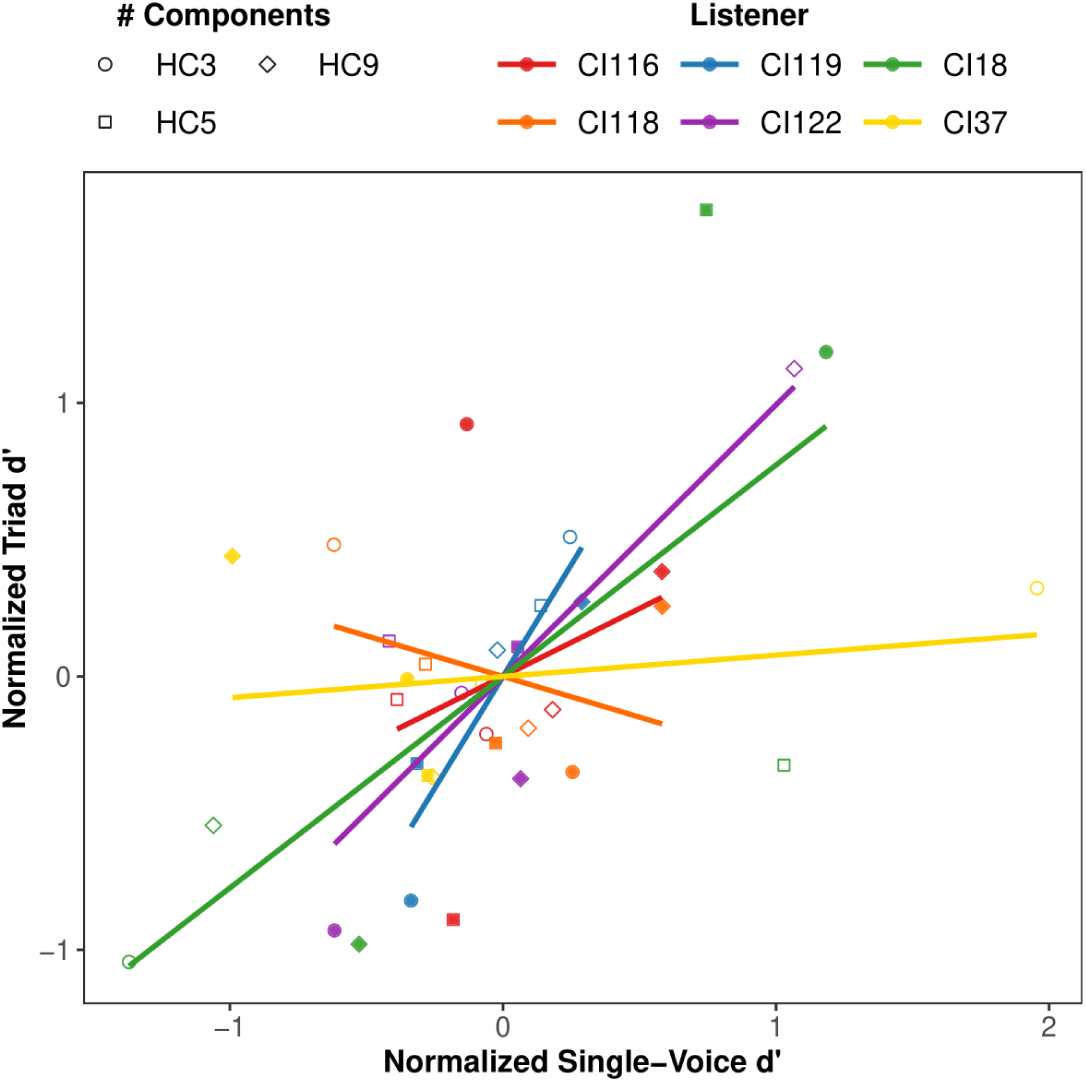
Partial within-subjects correlation (CI listeners indicated by color) between single-voice and triad *d’* scores normalized to remove effects of number of components per voice (indicated by marker type) and F0 range/VST (“low” indicated by empty markers, “high” indicated by filled markers). The VST low-high condition was excluded from the analysis.

#### 4. Comparison between discrimination of simultaneous and sequential triads

To directly compare discrimination sensitivity between simultaneous and sequential triad conditions, and to control for a possible confounding voice duration effect when varying the DC of sequential triads, we compared discrimination of simultaneous triads with durations of either 400 ms or 80 ms to sequential triads of different DC levels. As CI listeners were not able to discriminate sequential HC9 triads and as there was no difference in sensitivity between VST high and low-high conditions, we conducted this analysis for HC3 triads only and on *d’* scores averaged across the two high-voice change conditions (i.e., high and low-high) – VST low conditions were excluded as they did not yield any sensitivity. The averaged HC3 scores are shown inFigure 9. A one-factor ART ANOVA with five triad conditions as levels (i.e., simultaneous 400 ms vs. sequential DC100 vs. sequential DC75 vs. sequential DC50 vs. simultaneous 80 ms) yielded a significant effect, *F*(4, 20) = 11.52, *p* < .001, η²_G_ = 0.210. Post-hoc *t*-tests revealed better performance for simultaneous 400-ms triads as compared to simultaneous 80-ms triads, *t*(20) = 4.24, *p* = .001, sequential DC100 triads, *t*(20) = 5.75, *p* < .001, sequential DC75 triads, *t*(20) = 4.40, *p* < .001, and sequential DC50 triads, *t* = 5.95, *p* < 001. Notably, sensitivity for simultaneous 80-ms triads did not differ from any sequential-triad condition (*p* ≥ .206).

**Figure 9.**
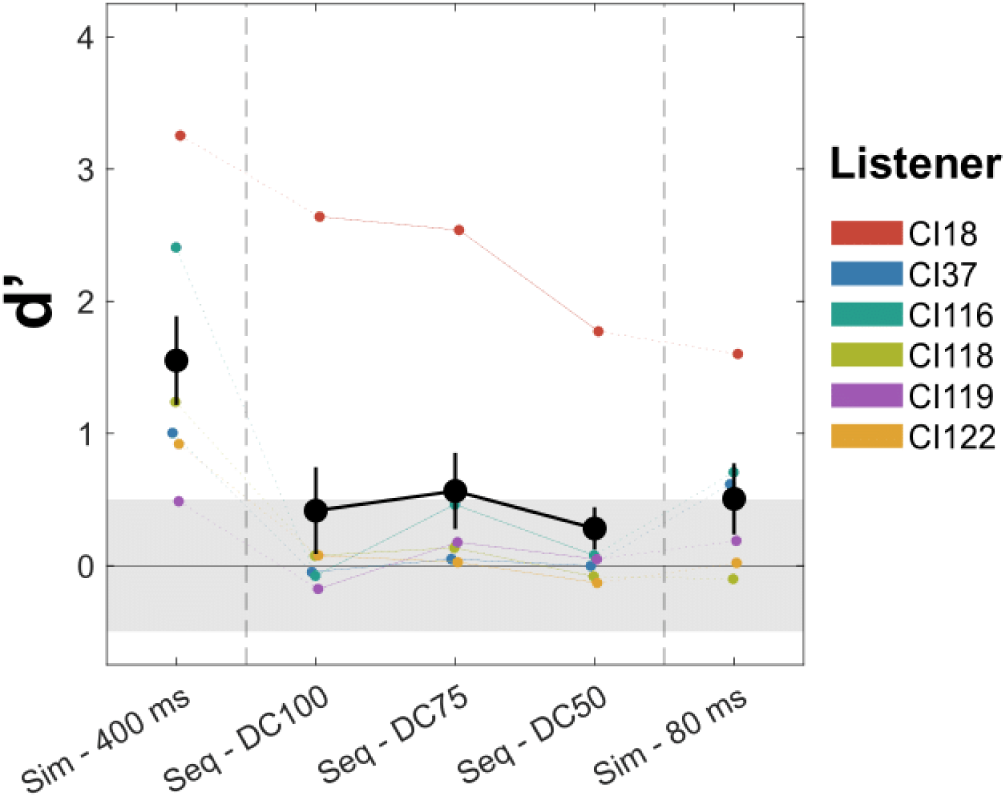
Discrimination sensitivity in terms of *d’* for selected **HC3** simultaneous and sequential-triad conditions. All other aspects as in Figure 6.

## III. SIMULATION: ENCODING OF TIMING AND PLACE OF EXCITATION IN AUTHENTIC FS4 PULSE PATTERNS

To better understand which cues were encoded by the CI processor with the CI listeners’ clinical fittings (crucially, filterbank configurations), we retrospectively analyzed the FS4 pulse patterns generated by the research processor for single-voice and simultaneous triad stimuli both with respect to temporal cues, that is, periodicities detected by the periodic-modulated Harmonic Locked Loop algorithm at the level of the individual electrode (PM-HLL; Hohmann, 2021), and place cues, that is, either the centroid of the CI-encoded “spectrum” (cf. Laneau et al., 2004; Swanson et al., 2019) or the maximum spectral contrast between the two triads to be discriminated in a condition of the psychophysical experiment. To this end, we converted the acoustic HC stimuli into electric multi-electrode pulse patterns using a research toolbox provided by MED-EL GmbH (Innsbruck, AUT). First, from the pulse patterns, we extracted coded periodicities and place-of-excitation information. Second, we attempted to relate those indicators with perceptual sensitivity. Importantly, we stress that this analysis does not capture all facets of electric stimulation but should rather be viewed as a means to assess which information may in principle be available to the CI listener and which information is already insufficiently coded at the electrode level.

Since we did not find perceptual sensitivity for sequentially presented triads, we included only single voices and simultaneously presented triads in the analysis.

### Methods

#### 1. Toolbox and stimuli

We replicated the electric pulse train stimuli that the CI listeners had to discriminate in the psychophysical experiment with a MATLAB implementation of MED-EL sound processors provided by MED-EL. This toolbox provides accurate emulation of the coding strategies and stimulation patterns as implemented in MED-EL sound processors, based on individual listener-specific clinical mapping parameters. Furthermore, we ensured that the acoustic amplitudes of the psychophysical experiment were accurately replicated in the simulations. Thus, we were able to closely simulate the listener-specific setup and electrode stimulation patterns for the psychophysical experiment, considering the individual differences between the CI listeners.

As “acoustic” input, we used only ‘different’ trials that were identical to those in the discrimination tasks. That is, each trial contained two stimuli and an inter-stimulus break. The first stimulus always contained the lower (voice) F0.

All simulations were conducted with MATLAB R2023b (The MathWorks, Inc., 2023).

#### 2. Extraction of timing and place-of-excitation information

Periodicities (and associated frequencies) were extracted from the pulse patterns elicited by each electrode in response to the input signal as an indicator of temporal coding. Periodicities were searched using a time-domain algorithm, the so-called Period-Modulated Harmonic Locked Loop (PM-HLL) algorithm (Hohmann, 2021). This algorithm has been shown to robustly estimate the F0 of acoustic signals with different types of periodicities and to track the F0s of the individual voices of simultaneous musical triads. It is entirely deterministic and not constrained to a specific type of temporal information such as, for example, fine structure or amplitude modulation. To make the “electric” pulse train stimuli better suitable for analysis, they were low-pass filtered using a sixth-order digital Butterworth filter with a cutoff frequency of 1000 Hz. The PM-HLL algorithm was run separately for each of the first six electrodes that were activated by the stimuli, of which electrodes one to four followed the FS coding approach (e.g., Hochmair et al., 2006; Müller et al., 2012) and electrodes five and six followed CIS coding (i.e., HDCIS; e.g., Müller et al., 2012). Starting with a predefined oscillator (search) frequency, the PM-HLL adapts and locks to a periodic signal component emerging in the input signal, if its fundamental frequency lies in a well-defined PM-HLL catch range. For the periodicity estimations reported here, the starting frequency was set at the geometric mean of the two targeted F0 or beat frequencies within a trial. The oscillator frequency requires a certain time to converge around a detected periodicity. Therefore, the first 60 ms of the search track were excluded from averaging to obtain a final periodicity estimate. For each detected periodicity, the standard deviation (SD) and the harmonic-to-noise ratio (HNR, in dB) of the estimate were calculated as indicators of periodicity saliency and estimation precision, respectively (Hohmann, 2021). From the periodicities (i.e., frequencies) estimated for the two stimuli of a discrimination trial, the FD was extracted in the form of a Weber fraction and converted to ST.

Place-of-excitation cues are only crudely coded with CIs (for a review, see e.g. McKay, 2005). Frequency components are hardly, if at all, resolved. Hence, spectral cues such as musical timbre are attributable to rather coarse, aggregate measures, for example, the spectral centroid (e.g., Macherey & Delpierre, 2013). With respect to pitch discrimination, such centroid shifts may well correlate with a perceptual “up-down” dimension related to changes in F0. However, since we employed a same/different task, other cues not necessarily corresponding to a change in F0 may have been used by the listeners for their responses. In an extreme scenario, listeners may have used simply any difference in spectral profile to perform the task. Such a cue may be well represented by the maximum spectral contrast between different stimuli, that is, the range of changes in electrode amplitude across all stimulating electrodes. Note, however, that CI listeners are not particularly sensitive to across-electrode changes in spectral profile. Instead, they perform within-electrode intensity analyses (e.g., Goupell et al., 2008).

In the pulse pattern analysis, we calculated per-electrode peak amplitudes as the averages (in *µ*A) of all pulses with amplitudes exceeding 90% of the maximum stimulus amplitude at the respective electrode. This was done to make the centroid estimate more robust to outlier pulses. To minimize residual effects of the automatic gain control, we omitted the first and last 100 ms from the stimuli. From the amplitudes, first, we calculated the spectral centroids (in mm, due to different electrode separations, cf. Table 1) of both stimuli of a discrimination trial and afterwards calculated the shift as the linear difference between centroids with electrode one being the 0-mm reference (cf. Laneau et al., 2004; Swanson et al., 2019). Second, we calculated the per-electrode changes in amplitude between first and the second stimulus, converted the differences to proportions of the electrode dynamic range (% DR), and calculated the peak contrast parameter as the range of amplitude changes, that is, the difference between maximum amplitude increase and maximum amplitude decrease (in % DR).

#### 3. Data processing and analysis

PM-HLL-estimated FDs were filtered so that only reasonably prominent and reliable estimates were included in the subsequent analyses. To this end, only FDs with a mean HNR > 0 dB and an SD at best 40 (single voices) or 20 (triads) times smaller than the mean F0 were included.

To exclude (quasi-)inactive electrodes from spectral analyses, only peak amplitudes estimated from more than four pulses (for each of the two stimuli, visualized in supplementary Figures S5 to S7) were considered reliable and used for the centroid shift and peak contrast estimates. The centroid shifts were calculated directly from the averaged peak amplitudes. The peak contrasts were calculated per electrode as the median peak amplitude difference derived from 1000 bootstrapped amplitude difference estimates.

To assess the fidelity of temporal encoding, within-subjects correlations *r* (Bland & Altman, 1994, 1995) between stimulus FDs and estimated FDs were calculated using R 4.5.1 (R Core Team, 2024) and its rmcorr package (Bakdash & Marusich, 2017, 2024). As with CIs no similar “ground truth” (i.e., a place analog to stimulus FD) exists for place-of-excitation cues, such analyses were not conducted for centroid shifts and peak contrasts. Still, to assess the relation between place-of-excitation cues and perceptual sensitivity, within-subjects correlations *r* were calculated between either centroid shift or peak contrast and the *d’* scores. Since the direction of centroid shift is irrelevant in a same/different task (as compared to, e.g., an up/down pitch discrimination task), we only included the absolute centroid shift in the analyses. Note that the peak contrast is an unsigned parameter by design.

To assess the relation between temporal cues and perceptual sensitivity, partial within-subjects correlations *r_p_* were computed between absolute^i^ estimated FD and *d’* scores using linear models and the emmeans package (Lenth et al., 2025). Since we did not consider across-electrode effects for temporal coding, the estimated FDs only incompletely represented the effect of the number of frequency components per voice.^ii^ To this end, the analyses factored out differential effects of the number of frequency components per voice by considering the marginal trend of estimated FD across HC3, HC5, and HC9 stimuli rather than the main effect of estimated FD. The trend was estimated from a linear regression model with *d’* as dependent variable. Included predictors were CI listener (categorical), number of frequency components per voice (categorical), and its interaction with estimated FD (continuous). The marginal trend (i.e., its slope) was then *t*-tested and the partial correlation *r_p_* was then derived from the *t* statistics (see Section II.A.4).

To jointly assess the relation between temporal cues, place-of-excitation cues, and perceptual cues, partial within-subjects correlations *r_p_* were calculated between absolute estimated FD, absolute centroid shift or peak contrast, and *d’* scores. An extended linear model was used that included absolute centroid shift or peak contrast as an additional continuous predictor. In all other aspects, the model was set up similarly to the above model for temporal cues only and *r_p_*’s were derived from the model *t* statistics. For all kinds of within-subjects correlations, 95% confidence intervals are reported in brackets.

### Results

#### 1. Single voices

Figure 10 shows the analysis results for single-voice stimuli. First, we investigated how precisely the FS4 strategy temporally encoded the F0s of the two voices to be discriminated in the same/different task and, in turn, how precisely the estimated FD followed the stimulus FD. The results are summarized in panel A for the low F0 range (F0s around 125 Hz) and in panel B for the high F0 range (F0s around 290 Hz). Across F0 ranges, the FS electrodes two to four encoded the stimulus FD with high precision and similarly well across number of frequency components per voice (i.e., number of components). FS electrode one, however, encoded the F0 less precisely. For the high F0, this was due to the F0s being largely outside the passband for almost all listeners. For the low F0s, this may be due to few HC frequency components generally falling into the passband. More importantly, however, the CIS electrodes five and six encoded the stimulus FDs with much lower precision, particularly in the high F0 range. We inspected the estimated F0s and confirmed that the PM-HLL did converge correctly, however, it seemed to detect periodicities related to the CIS carrier (e.g., the fourth carrier subharmonic closely falling together with the stimulus F0s in the high range) rather than the F0. As the carrier is constant across the two voices, the FD is expected to be close to zero in such a scenario.

**Figure 10.**
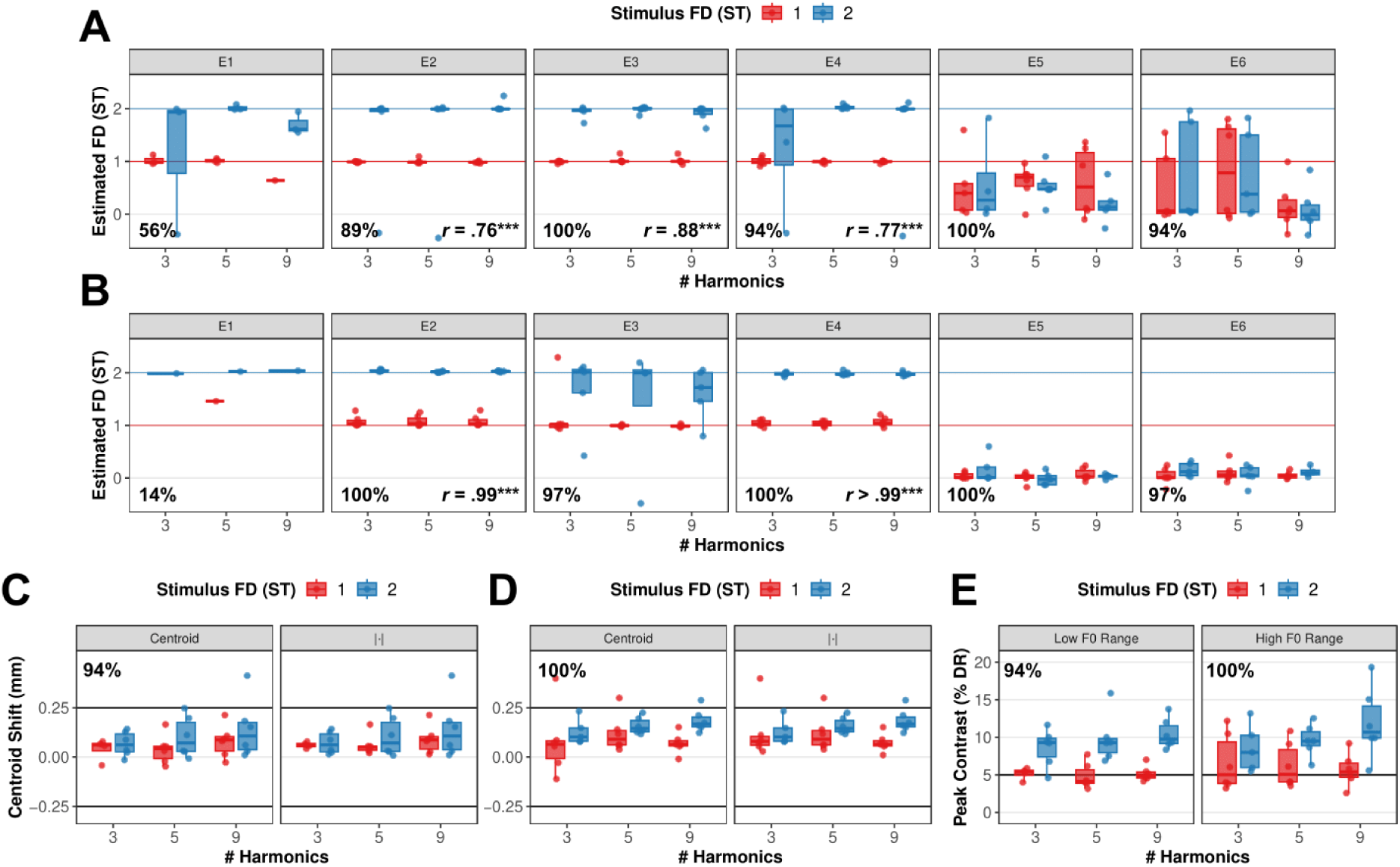
Results of the analysis of simulated FS4 pulse patterns with **single-voice stimuli** as acoustic input. **A** Frequency differences (FDs) in semitones (STs) estimated from pulse trains for the *low F0 range* at electrodes one to six (separate panels) using the Period-Modulated Harmonic Locked Loop algorithm (Hohmann, 2021) as a function of number of stimulus type (i.e., number of frequency components per voice; HC3, HC5, and HC9) and with stimulus FD as parameter. Lower left corners: rate of successful FD estimations. Lower right corners: point biserial within-subjects correlations of stimulus FD and estimated FD pooled across stimulus types (Bland & Altman, 1994, 1995). Only significant correlations are denoted. *… *p* < .05; **… *p* < .01; ***… *p* < .001. **B** Same analysis as panel A but for the *high F0 range*. **C** Estimated shift in place of excitation (i.e., spectral centroid in mm) for the *low F0 range*. Left panel: raw estimates; a positive shift indicates a genuine encoding of stimulus FD (i.e., an ascending F0, hence, a positive FD). Right panel: Absolute shift to reflect the fact that the psychophysically employed same-different task is insensitive to shift direction. Horizontal lines at ± 0.25 mm denote the smallest shift threshold reported in Laneau and Wouters (2004) and an indicator of absolute cue size. The proportion in the left panel indicates the share of successful estimates (see Section III.A.3 for details on data processing). **D** Same analysis as panel C but for the *high F0 range*. **E** Estimated peak contrasts (i.e., across-electrode range of changes in per-electrode peak amplitude between the two stimuli of a discrimination task trial) expressed as proportions of the dynamic range (% DR). The horizontal line at 5% DR indicates the threshold peak contrast estimated from the no-roving peak and notch detection thresholds reported in (Goupell et al., 2008). The proportions in the panels again indicate the share of successful estimates.

Next, we investigated the spectral centroid shift. Figure 9C shows the shifts for the low F0 range as a function of stimulus type with stimulus FD as parameter. Raw shifts (left panel) were small and largely fell below the smallest multi-channel place discrimination threshold of 0.25 mm as reported by (Laneau & Wouters, 2004). The analysis was set up such that basal (i.e., positive) shifts reflect the increase in F0 (i.e., the positive FD) from the first to the second single-voice stimulus in a discrimination trial. Hence, despite being small in size overall, shifts were consistent across listeners and, on average, seemed to encode the stimulus FD (i.e., the larger FD yielded larger shifts). Absolute shifts (right panel) were more similar across CI listeners than the raw shifts, thus, probably better representing differences between stimulus FDs. The difference in numbers of components was small, if present at all. Figure 9D shows the centroid shift estimates for the high F0. The pattern was similar to the low F0 range except that the stimulus FD was slightly better encoded.

Finally, we investigated the peak contrasts. Figure 10E shows the estimates for both F0 ranges in different panels. Across F0 ranges, contrast estimates were largely exceeding the thresholds reported for peak and notch detection by (Goupell et al., 2008). Furthermore, the stimulus FD was well reflected in the size of the peak contrast estimates. The number of components, however, did not seem to influence the peak contrasts. When comparing peak contrasts with centroid shifts, the effects of stimulus FD and number of components seemed to be largely similar for both measures, but the centroid shifts seemed to be smaller than thresholds reported in the literature while the peak contrasts seemed to be larger than peak/notch detection thresholds reported in the literature. When comparing both measures statistically, the within-subjects correlation was *r*(63) = .71 [.56, .81], *p* < .001, *N* = 70, underscoring that these measures were highly similar and mutually explained about one to two thirds of the variance.

Summarizing the single-voice analysis to this point, both the timing and the place of excitation may explain perceptual single-voice sensitivity. However, centroid shift estimates were overall small and potentially sub-threshold when compared to previous findings on place discrimination sensitivity. FS electrodes two to four temporally encoded F0 information with high acuity, but CIS electrodes five and six showed indications of distortion of temporal envelope information (cf. e.g., Goldsworthy et al., 2022). Thus, it may be the case that CI listeners extracted meaningful temporal information from FS electrodes that saliently encoded it. While such a conclusion assumes rather small effects of channel interactions (e.g., Balthasar et al., 2003) and favorable across-electrode timing (i.e., small delay between FS pulses across electrodes; e.g., de Groote et al., 2025; Lindenbeck et al., 2024), several studies showed decent temporal coding under such favorable conditions (e.g., de Groote et al., 2025; Griessner et al., 2021; Lindenbeck et al., 2024; Macherey & Carlyon, 2010). Furthermore, de Groote et al. (2025) found no evidence of across-electrode integration of information at apical electrodes with MED-EL CIs.

When quantitatively assessing the relation of temporal and/or place-of-excitation cues with perceptual sensitivity *in isolation*, (partial) within-subjects correlations with the *d’* scores were similar for absolute centroid shifts (*r*(63) = .44 [.22, .62], *p* < .001, *N* = 70), peak contrasts (*r*(63) = .52 [.32, .68], *p* < .001, *N* = 70), and absolute estimated FDs (*r_p_*(57) = .45 [.24, .61], *p* < .001, *N* = 68), explaining between 5 and 46% of the (residual) variance in perceptual sensitivity. When *jointly* assessing the relation between the *d’* scores, estimated FDs, and absolute centroid shift or peak contrast, both place-of-excitation cues correlated similarly with the *d’* scores (in both cases *r_p_*(54) = .34 [.09, .54], *p* = .010, *N* = 66) while the absolute estimated FD only correlated significantly when combined with the centroid shift (*r_p_*(54) = .34 [.09, .54], *p* = .010, *N* = 66) but not when combined with the peak contrast (*r_p_*(54) = .16 [-.11, .40], *p* = .248, *N* = 66).

Taken together, the analysis of simulated FS4 pulse patterns for single-voice stimuli suggests that place of excitation and/or timing at FS electrodes encoded stimulus F0. Hence, it may not have been necessary to elicit phase-locked neural firing to discriminate single-voice stimuli in the employed same/different task.

#### 2. Simultaneous triads

Figure 11 shows the analysis results for simultaneous-triad stimuli. To reduce the complexity of this analysis to a reasonable amount, we followed the “best”-electrode approach identified for the single voices and analyzed only FS electrode two. For the triad stimuli used in the psychophysical experiment, as summarized in the left table in panel A separately per VST, electrode two furthermore had the advantage that all three triad voices had only one voice frequency component in the listener-averaged electrode passband. For the middle and high voices, this was the F0, while for the low voice it was the first harmonic F1. Since filter slopes are typically shallow in CI filterbanks, the F0 of the low voice was also considered in the analysis below.

**Figure 11.**
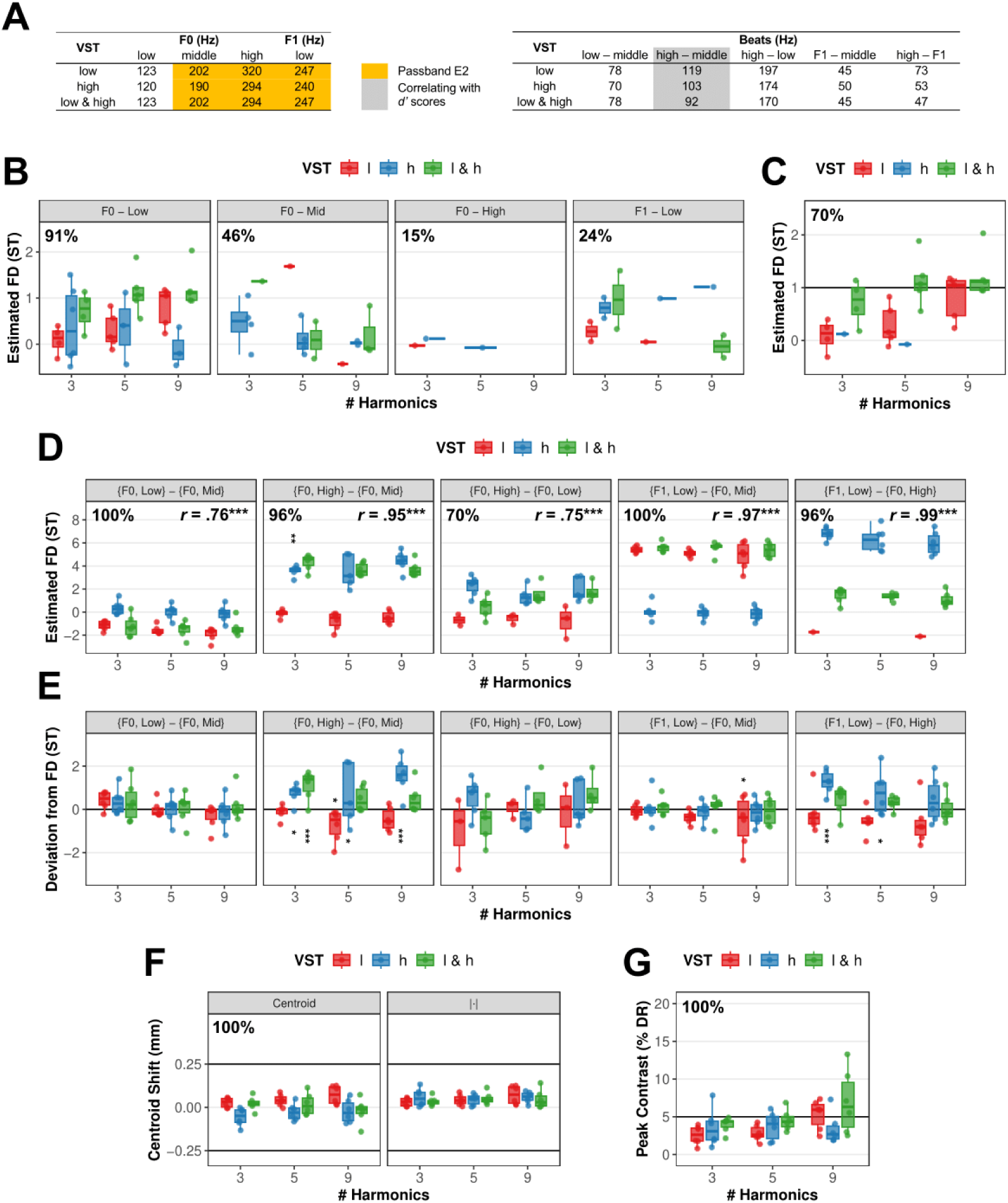
Results of the analysis of simulated FS4 pulse patterns for *electrode two only* and with **simultaneous-triad stimuli** as acoustic input. **A** Stimulus components at electrode two, most relevant in the pulse pattern analysis. Left: voice F0s and first harmonic F1 (low voice only) for the three VST conditions. Frequencies highlighted in orange fell within the *listener-averaged* passband of electrode two. Right: Geometric mean beat frequencies across the two triads presented in a same/different task trial (i.e., the two triads different by one ST in either one or two voices) separately per type of beat frequency. Beat type and mean frequencies highlighted in grey corresponded to perceptual sensitivity (i.e., the associated beat FDs correlated with the simultaneous-triad *d’* scores). **B** FDs estimated for voice F0s (low/mid/high) and the first harmonic F1 of the low voice as a function of number of frequency components per voice (# Harmonics) with VST as parameter. To match the stimulus FDs, estimates for low and high voices should have been 1 ST in case the VST matched the voice displayed in the respective panel (e.g., VST low matched F0 – low) and 0 ST otherwise. FDs for the middle voice should always be 0 ST. Upper left corners: share of successful FD estimations across # Harmonics and VSTs. Point biserial within-subjects correlations of VST FD and estimated FD (cf. Figure 10A) are not reported because they were never significant. **C** Estimated FDs for voice F0s rearranged from panel A so that only conditions with matching VST and voice F0 are displayed. Here, similarly to the 1-ST single-voice condition (cf. Figure 9A/B), the stimulus FD was always 1 ST, as indicated by the solid horizontal line. **D** Estimated FDs for *lower order voice difference frequencies* (i.e., beat FDs) instead of voice F0s. In each panel, the beat frequency is denoted at the top as a combination of two F0s/F1 previously investigated in panels B and C (cf. left table in panel A). The geometric mean frequencies of the beat frequencies occurring in the two triads of a discrimination task trial are summarized in the right table of panel E. Upper left corners: share of successful beat FD estimations. Upper right corners: point biserial within-subjects correlations of stimulus beat FD and estimated beat FD pooled across # Harmonics (cf. Fig. 9). Note the increased beat FD scale as compared to panels B and C. **E** As panel D, but for deviations (i.e., linear differences) of estimated beat FDs from stimulus beat FDs. Asterisks denote significant deviations from 0 as derived from bootstrapped linear-model marginal means (JASP 0.95.1; JASP Team, 2025). *… *p* < .05; **… *p* < .01; ***… *p* < .001. **F** Estimated spectral centroid shifts (in mm). All other aspects as in Figure 10C. **G** Estimated peak contrasts expressed as proportions of the dynamic range (% DR). All other aspects as in Figure 10D.

We first tested the influence of changed voice on triad discrimination (cf. e.g., Harrison & Pearce, 2020; Schneider & Wengenroth, 2009). To this end, Figure 10B shows the estimated triad voice FDs. Here, if VST and estimated voice match (e.g., VST low in panel “F0 – low” or VST low in panel “F1 – low”), the encoded FD should be 1 ST. Otherwise, it should be 0 ST (hence, for the “F0 – mid” panel, it should always be 0 ST as the middle voice was never changed). Across voices, it is evident that voice FDs were not well estimated. Furthermore, as indicated by the proportions in the top left corners, FD estimations failed regularly for all but the low-voice F0s. Figure 10B provides the subset of estimated voice FDs for which triad voice and VST matched. That is, here, VST-low and VST-high FDs should have been 1 ST while VST-low & high FDs should have been about 1.4 STs (simulating independent combination of low-voice and high-voice FD). Hence, here, the estimated FDs are visualized in a fashion similar to the single-voice FDs in Figure 11A/B. It again becomes obvious that voice F0 was not reliably encoded temporally, particularly for the VST high condition.

Next, to exploratively search for prominent periodicities (and associated frequencies) of *any* possible origin (e.g., F0, beating, subharmonics, etc.), we searched for the ten most prominent periodicities as determined by the HNR by scanning semi-tone bands across the entire F0-range (Hohmann, 2021). To reduce the bias of electrode, we conducted this search for all FS electrodes one to four. The results were qualitatively grouped and subsequently crudely attributed to a potential origin. The results of this analysis paralleled the voice-F0 analysis in that only a minority of extracted periodicities corresponded well with voice F0s (or their subharmonics). This exploratory analysis revealed that most detected periodicities potentially originated from beating between voice F0s (i.e., difference frequencies). Since beats are coded in the temporal envelope, irrespective of FS or CIS stimulation, we considered this result plausible and next analyzed only the simplest (i.e., lower order) beat frequency candidates to reasonably limit the scope of the analysis.

The beat frequencies included in the analysis are summarized in the right table of Figure 11A. Particularly, separately per VST, the geometric average beat frequency of the two triads of a discrimination trial are shown to provide an intuitive understanding of the modulation frequencies the analysis targeted. The beat frequencies themselves were linear differences between the voice F0s and F1 summarized in the left table of Figure 11A. Figure 11D shows the estimated beat FDs in separate panels per beat frequency type as a function of number of components and with VST as parameter. Based on the estimation-success rates in the top left panel corners, it is evident that all considered beats were detected much more reliably than voice F0s. Based on the correlations provided in the top right panel corners, it is furthermore evident that all of them closely resembled the stimulus beat FDs. Furthermore, it is evident that beat FDs tend to be larger than voice FDs (i.e., they were larger already in the stimulus, at least when expressed as Weber fractions), indicating that beat frequency changes might have been more easily detectable than voice F0 changes.

To assess whether the beat FDs were encoded genuinely (or whether the estimated beat FDs were, e.g., systematically elevated compared to the stimulus beat FD), we analyzed the deviations of the estimated beat FDs from the stimulus beat FDs. Figure 11E shows these deviations. While there were rare cases of significant deviations, these deviations did not seem to mimic the patterns in the behavioral data (cf. Figure 6). Thus, we concluded that beat frequencies are indeed genuinely coded in the pulse patterns of electrode two.

To explore whether any of the considered beats correlated with perceptual sensitivity in isolation (i.e., without place-of-excitation cues), we calculated partial within-subjects correlations between all five beat types (i.e., the estimated absolute beat FDs) displayed in Figure 11D and the simultaneous-triad *d’* scores. Indeed, the F0 high-F0 mid beat correlated significantly, *r_p_*(41) = .48 [.22, .65], *p* = .001, *N* = 52 (all other beats *p* ≥ .581).

Next, we investigated place-of-excitation information in the form of centroid shifts and peak contrasts. The shifts for simultaneous triads are summarized in Figure 11F. The structure of the analysis was identical to the single-voice analysis (cf. Figure 10C/D) except that VST rather than stimulus FD was the parameter. The results pattern, however, is different to that of the single-voice stimuli in that particularly the absolute shifts (right panel) were similar for all three VST conditions (while perceptual sensitivity was not, cf. Figure 6) and were generally barely measurable.

The contrasts for simultaneous triads are summarized in Figure 11G, again in similar fashion as for the single-voice stimuli (cf. Figure 10E). Also, here, the results differed from the single-voice analysis in that the contrasts were smaller---falling mostly below the threshold derived from (Goupell et al., 2008)---and furthermore seemed to be largest for the HC9 stimuli for which perceptual sensitivity was lowest (cf. Figure 6). Still, when comparing both measures statistically, the within-subjects correlation was *r*(47) = .52 [.28, .70], *p* < .001, *N* = 54, thus, mutually explaining about 8 to 49% of the variance.

To explore whether place-of-excitation cues correlated with perceptual sensitivity in isolation (i.e., without temporal beating cues), we calculated within-subjects correlations between either absolute centroid shifts or peak contrasts and simultaneous-triad *d’* scores. Neither shifts (*r*(47) = -.04 [-.32, .25], *p* = .797, *N* = 54) nor contrasts (*r*(47) = .07 [-.22, .34], *p* = .656, *N* = 54) correlated significantly with perceptual sensitivity, thus, further contrasting triad from single-voice results.

To assess whether temporal beating and spectral place-of-excitation cues correlated with perceptual sensitivity when considered *jointly*, we calculated partial within-subjects correlations between the triad *d’* scores, estimated absolute beat FDs, and absolute centroid shift or peak contrast. Contrary to the single-voice results, neither the absolute centroid shift (*r_p_*(40) = -.02 [-.32, .29], *p* = .896, *N* = 52) nor the peak contrast (*r_p_*(40) = .09 [- .23, .38], *p* = .586, *N* = 52) correlated significantly while the absolute beat FD correlatedsignificantly and similarly when combined with shift (*r_p_*(40) = .48 [.22, .64], *p* = .002, *N* = 52) or contrast (*r_p_*(40) = .48 [.22, .65], *p* = .001, *N* = 52).

Taken together, the analysis of simulated FS4 pulse patterns for simultaneous-triad stimuli suggests that temporally encoded difference-frequency (i.e., beat) information, at least at the examined electrode two, encodes musically relevant voice-F0 differences between triads rather than temporally encoded F0 information or spectrally encoded place-of excitation information in the form of centroid shifts or peak contrasts.

## IV. DISCUSSION

Due to technical, physical, and perceptual limitations of state-of-the-art CIs, listeners typically experience difficulties in perceiving pitch, especially in spectrally complex situations, for example, in the context of multiple concurrent pitches, similarly to speech in noise. Therefore, in the present study we investigated the influence of several stimulus parameters on triad discrimination in CI listeners. The aim of this investigation was to find favorable stimulus parameters which provide maximal perceptual salience of cues related to musical harmony. To that end, in a same/different triad discrimination task, we manipulated the spectral complexity (i.e., the number of frequency components) of triad voices, the triad voice in which a change occurred (i.e., the VST), and the temporal synchrony of triad voices. To vary the spectral complexity, triad voices were composed of either three-, five-, or nine-harmonic complex tones (HC3, HC5, or HC9 tones, respectively). We hypothesized that the lowest spectral complexity (i.e., HC3) yields the best performance. Regardless of the spectral complexity, a one-ST pitch change occurred either in the highest, the lowest, or both the highest and the lowest triad voices. We hypothesized that pitch changes in two voices (VST low-high condition) or in only the highest voice (VST high condition) yield better performance than a pitch change in only the lowest voice (VST low condition). To vary the amount of channel interactions between voice components, triads were presented either simultaneously (i.e., all triad voices were played at the same time) or sequentially (i.e., triad voices were played one after another). When triads were presented sequentially, we also manipulated the duty cycle (DC; 50, 75, or 100%) to control for potential effects of temporal masking. We hypothesized that sequentially presented triads yield better performance than simultaneously presented triads due to reduced spectral complexity and, thus, reduced interference between voices. We had no specific hypothesis regarding the DC as this variation was of an exploratory nature. Furthermore, to assess basic F0 sensitivity and to relate this sensitivity to triad perception, we also tested single-voice discrimination.

### Summary of findings

Our results showed that listeners were able to discriminate single voices and simultaneously presented triads but not sequentially presented triads. For simultaneous triads, in line with our hypotheses, discrimination was possible only for lower-complexity stimuli (i.e., HC3 and HC5) and for the VST high and low-high conditions. For sequentially presented triads, contrary to our hypothesis, listeners were insensitive for most of the tested conditions. When normalized for the effects of spectral complexity and F0 range, single-voice and simultaneous triad sensitivity was significantly correlated but this correlation explained at best about half the variance. Analyses of simulated FS4 pulse patterns revealed that single voices may have been discriminated based on both spectrally encoded place-of-excitation cues (i.e., centroid shift and peak contrast) and temporally encoded F0 cues. In contrast, simultaneous-triad sensitivity did not correlate with either place-of-excitation cue and voice F0s were only insufficiently coded. Instead, difference frequency (i.e., beating) cues could significantly explain some of the perceptual sensitivity.

The finding that beating cues may underlie simultaneous-triad discrimination sensitivity is in line with previous research showing exceptional beat frequency sensitivity in CI perception, even exceeding that of NH perception (Lu et al., 2014). Furthermore, triad perception solely based on temporal cues in the high frequency region (i.e., without any resolved harmonics, or place cue, being present) has been observed before in NH listeners (Graves & Oxenham, 2019). However, for HC9 stimuli, listeners were not sensitive despite the presence of beats. This may indicate a detrimental effect of carrier-modulator beats for CIS electrodes (Goldsworthy et al., 2022) interfering with the triad beats. It may also suggest that additional phenomena influenced triad perception such as, for example, across-electrode masking or channel interactions. In the following, the results are discussed in detail.

### Spectral complexity

Results revealed that sensitivity for harmonically relevant pitch differences of one ST within simultaneously presented triads improved with decreasing spectral complexity. These results confirmed our hypothesis that low spectral complexity is advantageous for CI-mediated perception of nuanced stimulus features such as small F0 changes within multiple concurrent tones. In the VST low condition, CI listeners could not discriminate triads, regardless of spectral complexity, revealing perceptual dominance of the high voice (consistent with our pilot test with NH listeners, see Supplementary A).

While previous studies investigating chord discrimination in CI listeners found low discrimination performance especially for natural sounds such as the piano (e.g., Brockmeier et al., 2011), our results show that spectral sparsity of stimuli might be one approach to counteract shortcomings of the CI such as extensive spread of electric fields and resulting spectro-temporal interactions. This is particularly relevant for studying chord discrimination, and, in further consequence, perception of musical harmony in general, as natural stimuli may confound results due to technical and physiological constraints of the CI rather than reflecting true perceptual capabilities.

Our finding that low spectral complexity is favorable for CI listeners is also in line with previous research reporting positive effects of spectral complexity reduction on melody perception and preference of musical stimuli (Galvin & Fu, 2011; Gauer et al., 2019; Nagathil et al., 2017; Nemer et al., 2017; for a review see Nogueira et al., 2019).

To obtain a broader view of the impact of spectral complexity, we investigated whether the single-voice discrimination data are predictive of the triad discrimination data. Single-voice sensitivity was present in all single-voice conditions, while performance was overall higher for the two-ST FD. Crucially, compared to previous investigations of single-voice discrimination with spectrally highly complex tones (e.g., piano tones; cf. Gfeller et al., 2002; Kang et al., 2009), here, using tones that were lower in spectral complexity, sensitivity was notably higher. However, while for simultaneous triads sensitivity was higher with three components compared to nine components per voice, the opposite was true for single voices. This might be due to overall differences in bandwidth (i.e., HC3 triad bandwidth was more similar to HC5 (low F0 range) or HC9 (high F0 range) single voices, cf. supplementary Figures S5 and S6). When comparing single voices and triads while controlling for the effects of F0 range/VST and spectral complexity, a significant residual within-subjects correlation remained. This correlation furthermore tended to be larger for musicians as compared to non-musicians. Combined with the results of the pulse pattern analysis, this trend may show that musicians were better at transferring their temporal-cue sensitivity from the F0 (for single voices) to difference frequencies (for simultaneous triads). However, since our experiment was not designed to test this conclusion, further targeted research is necessary to confirm (or reject) this trend.

### Changed voices and high-voice superiority

In addition to the spectral complexity hypothesis, our hypothesis regarding the changed voice or voices was confirmed as well. Listeners exhibited greater sensitivity when the change was in both high and low voices or when it was in the high voice only. This may mainly be explained by a perceptual bias for processing the highest voice more easily than lower voices which has also been observed before in NH listeners (Crawley et al., 2002; Fujioka et al., 2005; Palmer & Holleran, 1994) and was confirmed in our pilot test with a small NH listener cohort (Supplementary A). It was argued that this so-called high-voice superiority effect for natural tones originates in properties of the peripheral auditory system (Marie & Trainor, 2014; Trainor et al., 2014). Specifically, based on an established model of the auditory nerve (Ibrahim & Bruce, 2010; Zilany et al., 2009), the authors argue that the high-pass filtering of the middle ear and the lower cochlear gain for low frequencies attenuate low frequency components. Such effects may at least partly be restored by the high-frequency pre-emphasis filter employed in the front-end processing of the CI. Note, however, that while the mechanisms proposed by (Trainor et al., 2014) can only apply for simultaneously presented tones (see also Hove et al., 2014), interestingly, we observed a high-voice-superiority effect also for sequentially presented triads in our NH pilot tests, suggesting additional cognitive or learned mechanisms underlying this effect. The slightly larger sensitivity when both high and low voices changed compared to when only the high voice changed might have been observed simply because this condition comprised the largest amount of changing information.

Additionally, since the pulse pattern analysis has shown that beating between triad voices rather than voice F0s can partially explain the triad discrimination results, another potential reason for the insensitivity to changes in the low voice may be a perceptual dominance of a beat frequency that is not well associated with the low triad voice (i.e., it changes only when the high voice F0 changes). Beat frequencies between the high and the middle voice ranged from 92 to 119 Hz (see Fig. 9E). Given that CI listeners’ sensitivity to amplitude modulation is highest for modulation frequencies around 100 Hz (Chatterjee & Oberzut, 2011), this might partly explain the observed sensitivity for high-voice changes. However, since the relevant beat type (i.e., F0 high-F0 mid) explained at best about 40% of the variance in the *d’* scores (across all VST conditions), additional effects such as, for example, channel interactions as a function of spectral bandwidth, would need to be estimated and considered to explain more variance in the data. Still, we note that the frequency range of the relevant beat type (cf. right table in Figure 11A) was high enough to be largely outside the range of roughness perception as estimated for normal hearing (Plomp & Levelt, 1965). Other beat types falling into this roughness range were not correlated with perceptual sensitivity. Thus, both high-voice superiority and beating might contribute to explaining our results, however, to what extent each remains to be investigated.

### Sequential presentation of triad voices

When spectral complexity in our experiment was reduced by presenting triad voices sequentially instead of simultaneously, CI listeners’ performance dropped to chance level. Physical, perceptual, or cognitive challenges might have counteracted the theoretical advantage of reducing spectral complexity in sequential configuration.

First, the pulse pattern analysis suggested that CI listeners rely on beating between simultaneously presented triad voices rather than voice F0 information when discriminating simultaneous triads. Beating cues, however, are absent for sequentially presented triads, which likely at least partly explains the absent sensitivity to these triads. This is in line with the study by Lu et al. (2014), showing that an absence of beating cues eradicated CI listeners’ exceptional ability to frequency-tune guitar strings as compared to conditions with beating cues available. Furthermore, as here, place-of-excitation cues were partly predictive of sensitivity for single voices (but not for simultaneous-triad sensitivity) we might assume that, since sequential triads are brief single voices in succession, place-of-excitation cues should also underly sequential-triad sensitivity. However, the fact that sequential-triad sensitivity was absent supports the conclusion that, here, temporal pitch cues dominated spectral pitch cues for simultaneous triad perception. Second, even in sequentially presented single voices, spectral interactions between tone components might have been still too complex to be dealt with by the implanted auditory system within these short periods of time (i.e., 80-133-ms voice duration in sequential conditions as compared to 400-ms duration in simultaneous conditions). This is supported by the short (i.e., 80 ms) simultaneous condition for which performance dropped to chance level. Future investigations might consider longer voice durations in sequential triad discrimination; however, it is unclear whether sequential voices with longer durations would still be integrated into a harmonic percept and, thus, if this would measure harmony or rather melody perception.

Third, although we attempted to account for temporal masking effects by employing various duty cycles, excessive forward and/or backward masking might have impaired perception of sequential voices for CI listeners. Chatterjee & Kulkarni (2017) reported a mean estimated time constant of 99 ms, depending on stimulation electrode and masker level. By comparison, Oxenham & Moore (1994) reported time constants of <30 ms for NH listeners. For our sequential stimuli, thus, masking might have occurred particularly for DCs of 100% and 75% where tone durations were 133 ms or 109 ms and silence durations were 0 or 36 ms, respectively. Even for the shortest duty cycle of 50%, where tone and silence durations were both 80 ms, some amount of masking might have remained.

Fourth, sequential presentation might have caused higher listening effort compared to simultaneous presentation due to faster-changing auditory input. Higher listening effort has been associated with greater cognitive processing, loss of attention and withdrawal from the task (Marsella et al., 2017; Sherafati et al., 2022). Additionally, degraded auditory input, as also occurs with CIs, has been associated with impaired memory for what has been heard (despite the degraded input remaining intelligible; e.g., Ward et al., 2016) and even impaired memory for non-degraded input adjacent to the degraded input (Cousins et al., 2014; Piquado et al., 2010). This memory impairment may have further exacerbated the working memory challenges inherently present in the sequential condition.

Interestingly, one CI listener (CI18) in our sample exhibited extraordinarily high sensitivity in the sequential triad conditions, leading to the idea that the low average performance in this condition is not due to some CI-inherent limitation that is impossible to overcome. We can only speculate why CI18 was the only participant being able to perform at such a high level in this condition. Yet, it is important to note that CI18 actively plays several musical instruments since childhood, which is known to positively influence perceptual abilities and cognitive functioning (e.g., Parbery-Clark et al., 2011). Furthermore, he has no residual hearing in both ears, while the other musicians in our sample have residual hearing (through hearing aids) on the contralateral ear, and both confirmed that in everyday listening situations, especially when it comes to playing music, they rely more on the hearing aid ear than on the CI ear.

### Limitations and suggestions for future investigations

As we tested a small, relatively homogenous sample, it would be informative to replicate the experiment with a larger number of participants including different implant types, stimulation strategies, and etiologies. Also, despite the employed loudness adjustment, some possibility of subtle pitch-induced loudness changes contributing to discrimination performance might have remained. We argue, however, that the exclusive use of loudness cues would not explain decreasing performance with increasing number of components as there is no reason to assume less useful loudness cues for tones with larger numbers of components. Roving levels of individual triad voices would be one approach to further reduce listeners potential reliance on loudness cues. However, as level roving induces changes of timbre, this might render the task even more difficult for the listeners. Finally, while our investigations only considered basic peripheral discrimination of complex-tone triads (which is, however, the fundamental basis of broader musical harmony perception), future research should explore higher-order harmonic processing such as, for example, processing of multiple-chord harmony progressions with spectrally reduced tones.

## V. CONCLUSIONS

We showed that CI listeners were able to discriminate harmonically relevant pitch changes of one semitone within complex-tone triads (i.e., three simultaneously presented tones) when spectral complexity of triad voices was largely reduced and when the pitch change was either in both the high and low voices or only in the high voice, but not when it was in the low voice only. Specifically, performance was higher for three-component compared to nine-component harmonic complex tones. When triad voices were presented sequentially to reduce the potential for spectral interference in CI stimulation, average CI performance dropped to chance level. Still, CI listeners were on average able to discriminate single voices with a frequency difference of one semitone. Furthermore, single-voice discrimination sensitivity predicted simultaneous-triad discrimination sensitivity when controlling for spectral complexity and changed voice or F0 range, respectively. With respect to sequentially presented triads, we suggest that physical, perceptual, and cognitive factors such as voice duration, lack of beating between triad voices, temporal masking, or cognitive load might have contributed to the degraded sensitivity compared to the simultaneous condition.

Our analysis of electrical stimulation patterns suggests that the discrimination of simultaneous triads was largely based on temporal cues, despite the discrimination of single voices being based on temporal and place-of-excitation cues (if not largely based on the latter). Still, a multitude of effects too differential to be examined with the current sample of CI listeners such as auditory health, channel interactions, and across-electrode integration of information prevent a definite conclusion at this stage.

While future investigations might attempt to clarify the lack of discrimination sensitivity for sequential triads, our main finding, that is, improvement of simultaneous-triad discrimination by reduction of spectral complexity, has direct implications for future investigations towards more specific musical relevance. In particular, we suggest examining higher-level, cognitive perception of complex musical features such as consonance versus dissonance or harmonic syntax using stimuli tailored to the particular shortcomings of current CI systems.

## VI. SUPPLEMENTARY MATERIAL

See supplementary material A for normal hearing listeners’ data, supplementary material B for selected questions from the GOLD-MSI (Schaal et al., 2014), and supplementary material C for additional figures of the CI pulse pattern simulation.

## Author declarations

### Author contributions

MA and ML share first authorship. All authors contributed to concept and study design. MA programmed the experimental procedure while ML programmed the stimuli and CI processor. ML mainly contributed to data acquisition and performed the main statistical analyses. MA performed the post-hoc testing and wrote the first draft of the manuscript except the Methods section. ML wrote the Methods section and created the figures. ML and BL conducted the pulse pattern analysis. All authors revised and proofread the manuscript.

## Acknowledgments

We thank our listeners for their patience and passion. We thank Josh Stohl, Reinhold Schatzer, Dirk Meister, and Peter Nopp for providing the toolbox to simulate the research processor.

## Ethical considerations

In the experiments, we followed the European Charter of Fundamental Rights, worked along the guidelines of “Good Scientific Practice,” and fulfilled the ethical principles for research involving human subjects. The study was conducted in accordance with the ethics statement for human experimental research conducted at the Acoustics Research Institute of the Austrian Academy of Sciences, as approved by the Commission for Science Ethics of the Austrian Academy of Sciences on April 23, 2024.

## Data availability

The datasets generated during and/or analyzed during the current study are available from the corresponding author on reasonable request.

## Footnotes

i Similarly to the centroid shift, the direction of estimated rate change is irrelevant in a same/different task.

ii For example, an increase in frequency components may result in more components falling into the frequency band assigned to a certain electrode which in turn affects both the timing (FS coding only) and the amplitude of the electrode’s pulses (both FS and CIS coding). However, this effect 1) may differ between electrodes and 2) may be rather rare as most electrodes will always encode at least a few acoustic frequencies due to their broad tuning. Beyond that, any detrimental effect of channel interactions which are assumed to be more severe for broadband stimuli are not captured at all by the present analysis of temporal encoding.

