## Supplementary A, B, c for "Discrimination of spectrally sparse complex-tone triads in cochlear implant listeners"

### Single-Voice and Triad Discrimination in Noise by Normal-Hearing Listeners

#### Methods

To qualitatively justify our hypotheses and verify that particularly sequential triads can in principle be discriminated with the planned experimental setup, four normal-hearing (NH) listeners (2 females, 2 males, $M[SD]_{age}$ = 28[2.8] years), for which age-adequate normal hearing was confirmed with audiograms, participated in pilot tests. Three listeners had at least amateur musical training. Two of the listeners were the first and second author, respectively. Listeners were always tested with their right ear only.

The stimuli and experimental setup were identical to the CI experiment. The acoustic signal was presented to the right ear using circumaural audiometric Sennheiser HDA 200 headphones (Sennheiser electronic GmbH & Co. KG, Wedemark, GER) at a sound pressure level of 56 dB(A). Furthermore, to mask combination tones and to approximately match NH with CI performance in the main experiment (i.e., discrimination of simultaneously presented triads, see Section II.B.2.a) and Figure 6), white noise was added at a signal-to-noise ratio (SNR) of -6 dB such that the measured total sound pressure level was 63 dB(A). Although we are aware that adding white noise is not optimal for reducing pitch resolution in NH listeners (cf. Gockel et al., 2006), we considered this the most feasible way to avoid ceiling effects and, therefore, non-interpretability of results. Importantly, our goal was *not* to simulate CI performance in NH listeners. The white noise was digitally generated and bandpass filtered between 30 and 3300 Hz using Pure Data 0.49.0 (www.puredata.info). It was presented continuously only to the right ear. All listeners were tested in an anechoic chamber (IAC 1202-A, IAC GmbH, Mönchengladbach, GER). No loudness balancing was performed, but all stimuli (triads and single voices) were adjusted to have the same overall root-mean-square (RMS) amplitude. This means that for simultaneous triads the individual voices had lower RMS amplitudes compared to the single-voice stimuli.

To statistically analyze the NH d’ scores, we used ART ANOVAs (Wobbrock et al., 2011) similarly to the CI data.

#### Results

##### *Single-voice discrimination*

Sensitivity (in terms of *d’* alongside normalized [within-subjects] standard errors; Cousineau, 2005; Morey, 2008) is shown in Figure S1 as a function of the number of components with frequency difference (FD) as parameter and in separate panels for low and high F0 Range, respectively. As to be expected, group sensitivity saturated and, hence, no statistical analyses were performed.

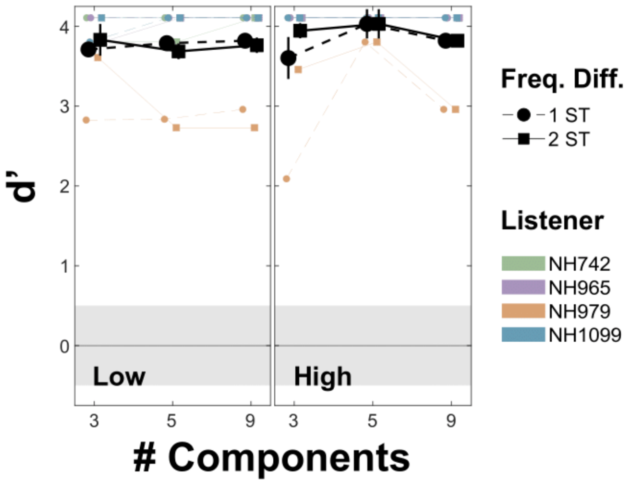


*Figure S1. Discrimination sensitivity in terms of d' for single voices as function of frequency components per voice. Panels distinguish F0 ranges (low vs. high). Marker shapes and line types denote the frequency difference (FD) to be discriminated in terms of STs. Error bars denote normalized (i.e., within-subjects) standard errors. The grey ribbons indicate chance performance for 100 repetitions. Individual data in color.*

*Discrimination of simultaneously presented triads*

Figure S2 shows discrimination sensitivity for simultaneous triads as a function of number of components with changed voice(s) as parameter. On the group level, sensitivity decreased with increasing number of components and was highest for low & high-voice changes and lowest for low-voice changes. NH listeners furthermore showed a pattern broadly similar to CI listeners (cf. Figure 6), although the NH experiment did not attempt to simulate CI listening.

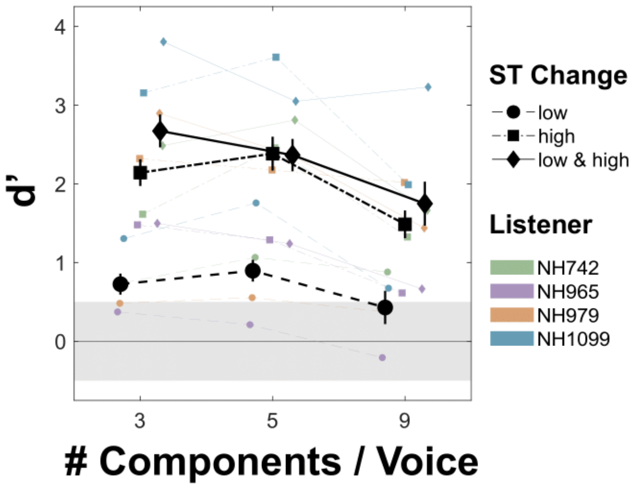


*Figure S2. Discrimination sensitivity in terms of d' for simultaneous triads as function of frequency components per triad voices. Marker shapes and line types denote the triad voice(s) with a ST change to be discriminated. All other aspects as in Figure S1.*

An ART ANOVA revealed a main effect of changed voice(s), $F(2, 6) = 48.30$, $p < .001$, $\eta^2_G = 0.510$, due to higher sensitivity for high-voice than low-voice changes, $t(6) = 7.41$, $p < .001$, as well as higher sensitivity for low & high-voice- than low-voice changes, $t(6) = 8.64$, $p < .001$. Furthermore, we found a main effect of number of components, $F(2, 6) = 12.45$, $p = .007$, $\eta^2_G = 0.167$, due to higher sensitivity for HC3 triads than for HC9 triads, $t(6) = 4.90$, $p = .004$, as well as higher sensitivity for HC5 triads than for HC9 triads, $t(6) = 4.94$, $p = .004$.

For NH listeners, we did not run bootstrapped *t*-tests against zero as absolute NH performance was fully dependent on the specific SNR we chose.

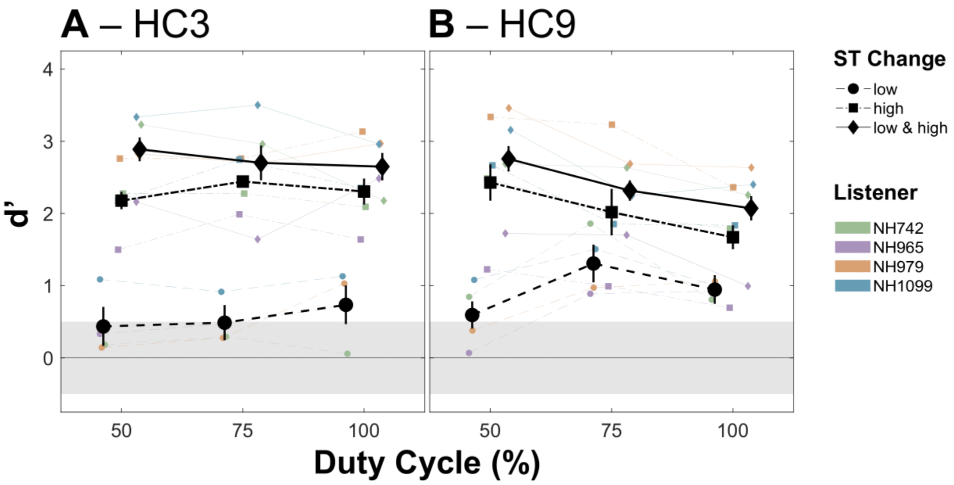


*Figure S3. Discrimination sensitivity in terms of d' for sequential triads (**A** HC3 and **B** HC9) as a function of duty cycle (DC). All other aspects as in Figure S2.*

*Discrimination of sequentially presented triads*

Figure S3 shows discrimination sensitivity for sequential triads for HC3 triads (panel A) and for HC9 triads (panel B) as a function of duty cycle (DC) with changed voice(s) as parameter. On the group level, sensitivity was high for high-voice- and low & high-voice changes while for low-voice changes it was lower with HC9 triads and mostly absent with HC3 triads. Furthermore, for HC9 triads in high-voice or low & high-voice conditions, sensitivity decreased with increasing DC.

An ART ANOVA for the HC3 *d'* scores revealed a significant main effect of changed voice, $F(2, 6) = 23.29$, $p = .001$, $\eta^2_G = 0.820$. The main effect of DC, $F(2, 6) = 0.12$, $p = .886$, $\eta^2_G = 0.003$ and the interaction, $F(4, 12) = 2.77$, $p = .076$, $\eta^2_G = 0.059$, were not significant. An ART ANOVA for the HC9 *d'* scores revealed a significant main effect of changed voice, $F(2, 6) = 13.91$, $p = .006$, $\eta^2_G = 0.537$. The main effect of DC was not significant, $F(2, 6) = 3.37$, $p = .104$, $\eta^2_G = 0.075$, but the interaction was significant, $F(4, 12) = 6.34$, $p = .006$, $\eta^2_G = 0.159$. We again did not run bootstrapped *t*-tests against zero.

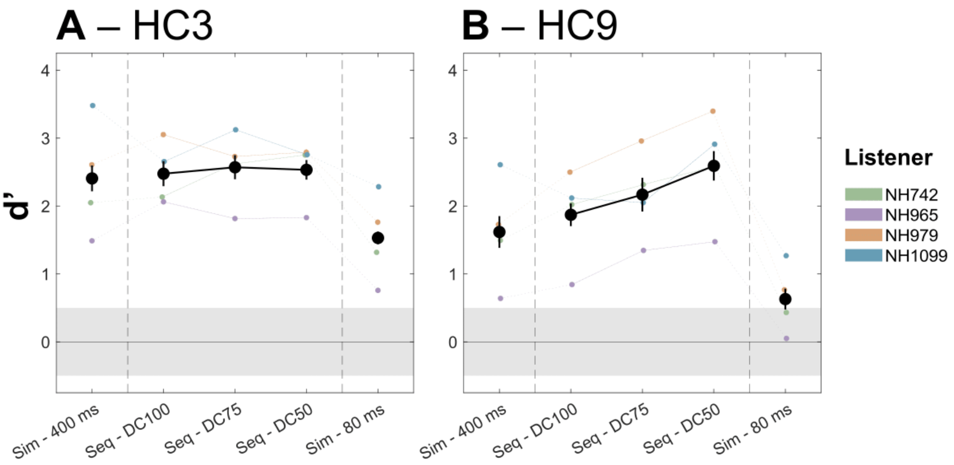


*Figure S4. Discrimination sensitivity in terms of d' for selected triad conditions in separate panels for **A** HC3 and **B** HC9. All other aspects as in Figure S2.*

*Comparison between discrimination of simultaneous and sequential triads*

To assess whether sequential triads can in principle be discriminated and to estimate to what extent the voice duration effect confounds the DC manipulation, we compared discrimination of simultaneous triads with durations of either 400 ms or 80 ms and sequential triads of different DC levels. As there was little sensitivity for low-voice change conditions, we conducted this analysis on *d′* scores averaged across the two high-voice change conditions (i.e., high and low & high). The *d′* scores are shown in Figure S4 for HC3 (panel A) and HC9 (panel B) stimuli.

For HC3, the one-factor ART ANOVA revealed a significant effect, $F(4, 12) = 6.70$, $p = .005$, $\eta^2_G = 0.325$, due to lower scores for simultaneous 80-ms triads as compared to simultaneous 400-ms triads, $t(12) = -3.47$, $p = .012$, sequential DC100 triads, $t(12) = -4.09$, $p = .005$, sequential DC75 triads, $t(12) = -4.34$, $p = .005$, and sequential DC50 triads, $t(12) = -4.20$, $p = .005$. Sensitivity for simultaneous 400-ms triads did not differ from any sequential-triad condition ($p = .790$). For HC9, the one-factor ART ANOVA revealed a significant effect, $F(4, 12) = 16.30$, $p < .001$, $\eta^2_G = 0.503$. Post-hoc *t*-tests revealed performance for simultaneous 400-ms triads to be higher than for simultaneous 80-ms triads, $t(12) = 3.64$, $p = .007$, lower than for sequential DC50 triads, $t(12) = -3.93$, $p = .005$, and marginally lower than for sequential DC75 triads, $t(12) = -2.32$, $p = .056$. Furthermore, performance was lower for simultaneous 80-ms triads than for sequential DC100 triads, $t(12) = -4.85$, $p = .001$, sequential DC75 triads, $t(12) = -5.96$, $p < .001$, and sequential DC50 triads, $t(12) = -7.57$, $p < .001$.

#### Conclusion

As expected, NH listeners performed exceptionally well in single-voice discrimination with such simple stimuli. For triads, NH listeners could discriminate simultaneous and sequential triads similarly well. They generally favored low spectral complexity and changes in the high as compared to changes in the low triad voice. For sequential triads, shortening the DC (and thereby the voice duration) had either no (HC3) or a beneficial (HC9) effect. For simultaneous triads, however, shortening the stimulus duration (and thereby the voice duration) worsened discrimination performance, highlighting a potential connection between spectral complexity and stimulus duration.

In summary, the NH pilot data largely confirmed our expectations, and we therefore considered our experimental setup feasible to test the hypothesized effects with CI listeners.

#### Supplementary B:

#### Music Questionnaire

Selected questions from the Goldsmiths Musical Sophistication Index (Gold-MSI; Müllensiefen et al., 2014). Participants were presented with German translations of the questions (Schaal et al., 2014). Questions 1 and 2 were answered using a rating scale (completely disagree – strongly disagree – disagree – neither agree nor disagree – agree – strongly agree – completely agree). Questions 3-5 were answered using pre-defined options for numbers of years (see below).

1. I can tell when people sing or play out of tune.
2. I would not consider myself a musician.
3. I have had *0 / 0.5 / 1 / 2 / 3-5 / 6-9 / 10 or more* years of formal training on a musical instrument (including voice) during my lifetime.
4. I have had formal training in music theory for *0 / 0.5 / 1 / 2 / 3 / 4-6 / 7 or more* years.
5. I can play *0 / 1 / 2 / 3 / 4 / 5 / 6 or more* musical instruments.

#### Supplementary C:

#### Simulations of FS4 Pulse Patterns – Figure Captions

Figure S5. Estimated peak amplitudes of single-voice stimuli as proportions of the dynamic range (% DR) as a function of electrode and with stimulus # ($1^{st}$ stimulus with lower F0: solid line; $2^{nd}$ stimulus with higher F0: dashed line) as parameter. Columns distinguish frequency components per voice (i.e., HC3, HC5, and HC9) and rows distinguish F0 differences (FD, in semitones). **A** Low F0 range. **B** High F0 range.

Figure S6. Estimated peak amplitudes of simultaneous-triad stimuli expressed in % DR as a function of electrode and with stimulus # ($1^{st}$ stimulus with lower F0: solid line; $2^{nd}$ stimulus with higher F0: dashed line) as parameter. Columns distinguish frequency components per voice and rows distinguish VST conditions.

Figure S7. Estimated peak amplitude differences (in % DR) as a function of. Columns distinguish frequency components per voice. **A** Single-voice stimuli. FD is the parameter and rows distinguish F0 ranges. For each combination of combination of frequency component per voice, F0 range, and FD, the peak contrast defined is defined as the range of amplitude differences across electrodes. **B** Simultaneous-triad stimuli. VST is the parameter. For each combination of frequency component per voice and VST, the peak contrast is defined as the range of amplitude difference across electrodes.
