## Supplementary figures and images for "Discrimination of spectrally sparse complex-tone triads in cochlear implant listeners"

### Supplemental Figure 5

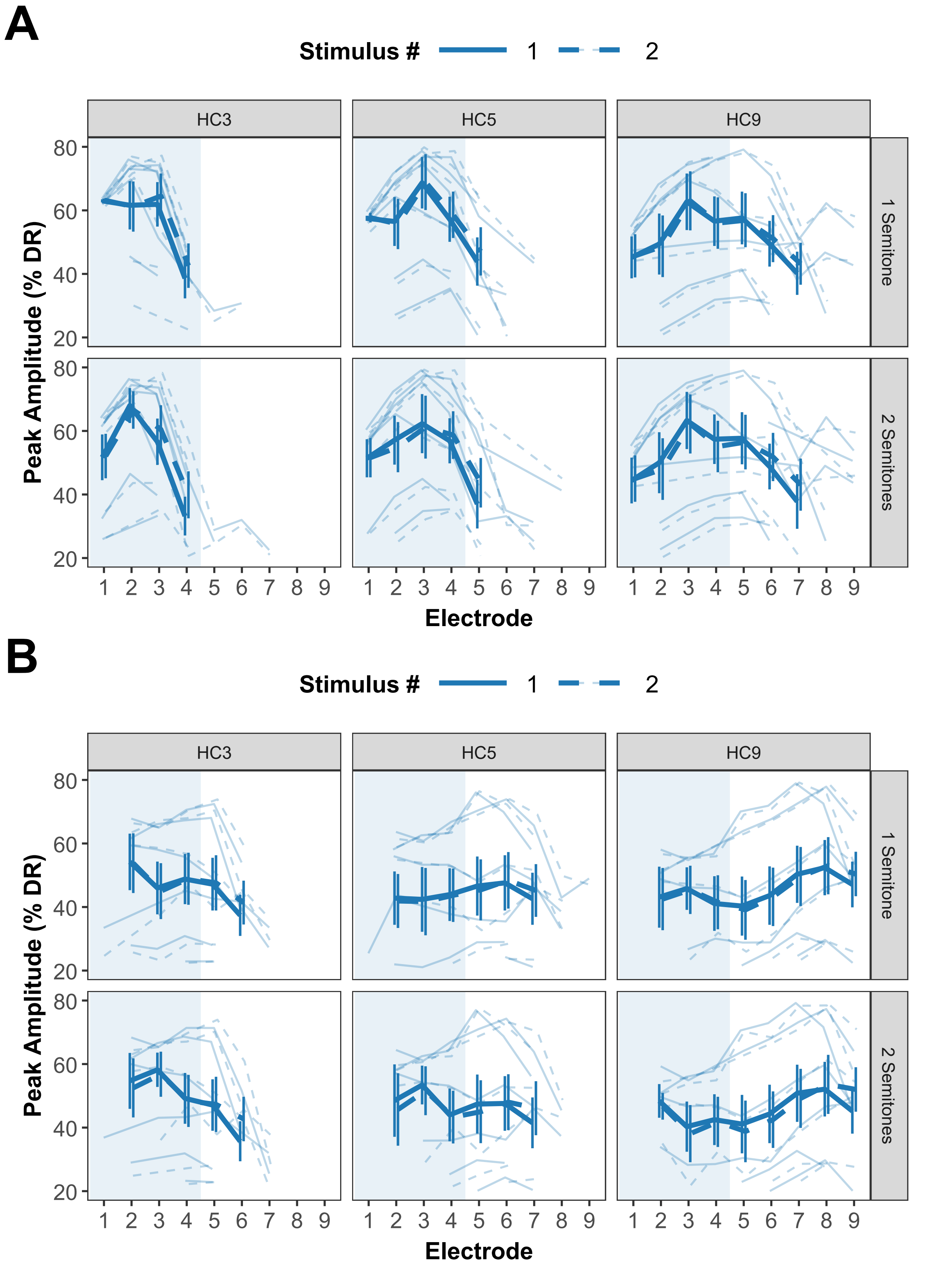

### Supplemental Figure 6

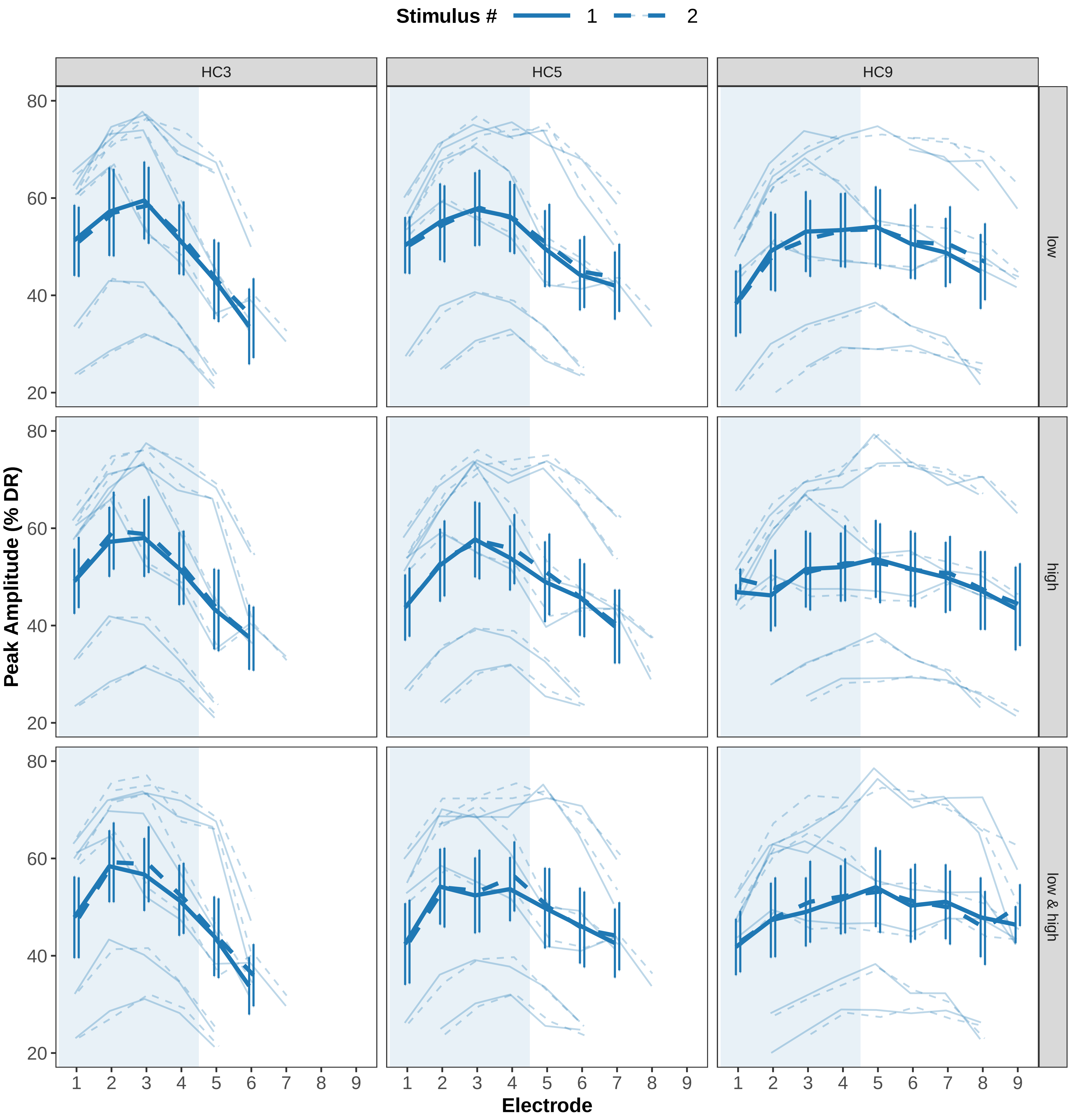

### Supplemental Figure 7

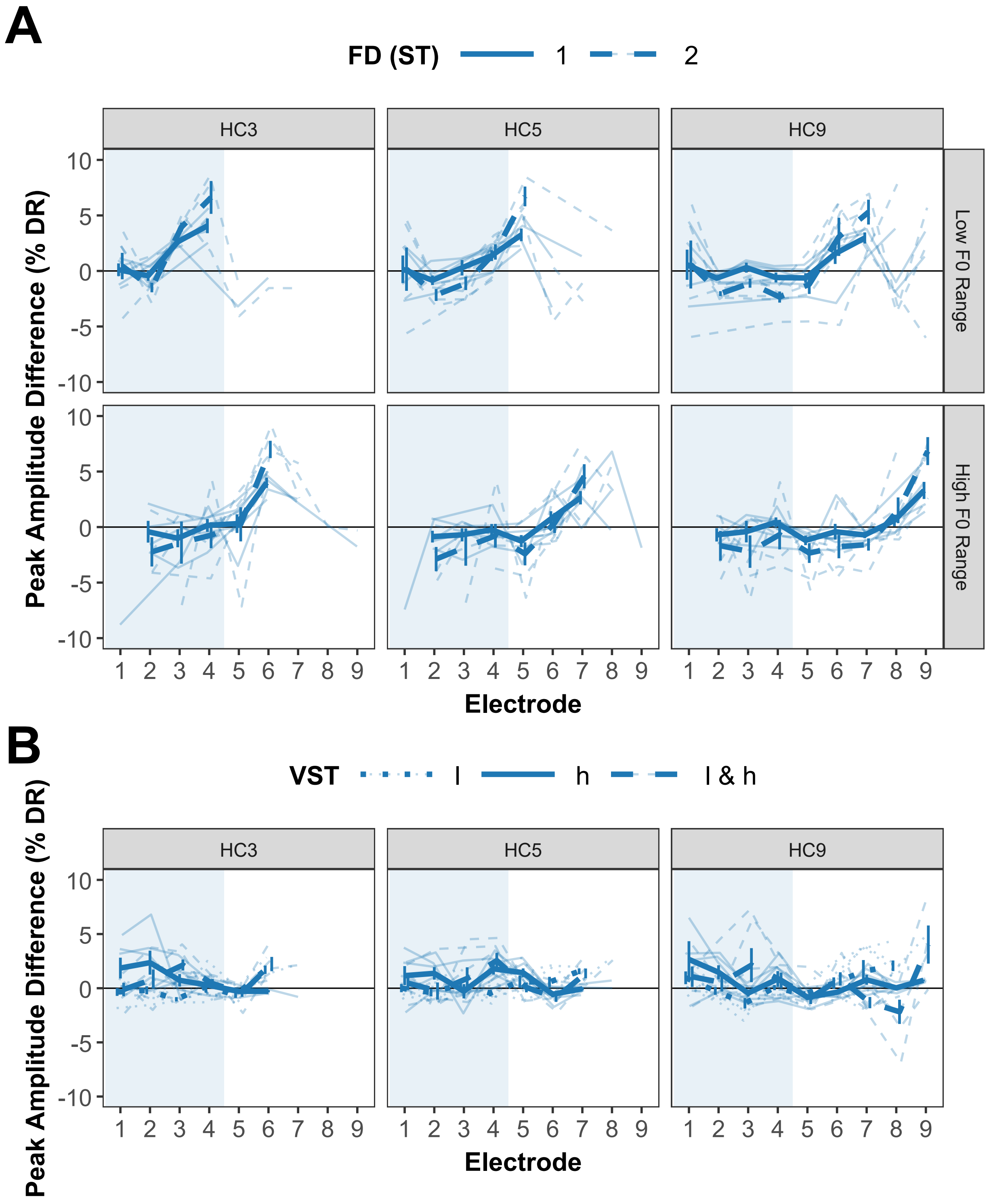
